# Quantitative assessment of sea slugs (Heterobranchia) assemblages along the western coast of Kyushu, Japan: a baseline for long-term biodiversity monitoring

**DOI:** 10.64898/2026.01.30.702737

**Authors:** Riko Kato, Mitsuharu Yagi

## Abstract

Quantitative information on the seasonal dynamics of heterobranch sea slug assemblages remains limited in warm-temperate coastal regions, despite their ecological importance as benthic consumers and indicators of environmental change. Here, we conducted a standardized, multi-seasonal SCUBA-based survey of sea slug assemblages at two rocky reef sites (Tatsunokuchi and Nomozaki–Akase) along the northwestern coast of Kyushu, Japan, from February 2024 to November 2025. Across the study period, a total of 81 species comprising 892 individuals were recorded. Species richness and total abundance exhibited pronounced seasonal variation at both sites, with higher values in winter–spring and marked declines during summer. Assemblage composition shifted seasonally from relatively even communities in winter–spring to dominance by a few taxa in summer, a pattern reflected by concurrent changes in diversity indices. Water temperature displayed clear seasonal cycles and was negatively correlated with both species richness and total abundance, indicating a close association between thermal conditions and seasonal changes in sea slug assemblages. While causal mechanisms were not explicitly tested, these consistent patterns highlight the importance of temporal environmental variability in structuring heterobranch communities in this region. This study provides one of the few quantitative, multi-seasonal baselines of heterobranch sea slug assemblages in warm-temperate coastal Japan, offering a reference framework for future ecological monitoring and assessments of environmental change.

## INTRODUCTION

Climate change–driven alterations in ocean conditions are restructuring coastal marine communities worldwide (Poloczanska et al., 2013; Pinsky et al., 2020). Rising seawater temperatures have accelerated the poleward expansion of warm-affinity species and the contraction of cold-affinity taxa, a phenomenon widely recognized as *community tropicalization* (Vergés et al., 2014; Wernberg et al., 2016). Around Japan, one of the world’s fastest-warming ocean regions, similar shifts have been documented across a broad range of coastal assemblages (Hongo & Yamano, 2013; Ota et al., 2021), highlighting the urgent need to understand how warming alters biodiversity patterns and community dominance structures.

Long-term and standardized monitoring is essential for quantifying ecological responses to climate change (Rees et al., 2007; Yasuhara et al., 2020). Sea slugs (Heterobranchia) occur across tropical to cold-temperate regions and play ecologically important roles through their feeding interactions with sponges, algae, and bryozoans (Gosliner et al., 2008; Nakano, 2019). Because many species exhibit strong sensitivity to temperature and habitat conditions, shifts in sea slug distributions and assemblage structures have been reported in response to environmental change, positioning them as promising indicators for detecting climate-driven biodiversity shifts (Nimbs & Smith, 2015; Chou et al., 2022).

Recent studies from western Kyushu, Japan have suggested multi-decadal changes in sea slug assemblages. A synthesis of six decades of records reported a marked increase in tropical species and a decline in cold-affinity taxa, indicating substantial long-term community restructuring (Kato & Yagi, 2025). However, most sea slug surveys in Japan—including historical datasets—are based on one-time visual observations lacking standardized quantitative methodology (Nakano, 2019; Kashio et al., 2021). As a result, comparisons among regions or across time are hindered by variation in survey effort and recording practices (Ota et al., 2021). Even the long-term dataset of Kato and Yagi (2025), although invaluable for revealing broad temporal trends, does not resolve fine-scale community structure at individual sites nor clarify how present-day assemblages relate to local microhabitat conditions. This represents a critical knowledge gap, as detecting early signals of tropicalization or community turnover requires robust, repeatable, site-level quantitative baselines.

To address this gap, we conducted standardized quantitative line-transect surveys at two coastal sites in western Kyushu (Tatsunokuchi and Akase, Nagasaki Prefecture, Japan) to characterize the present-day sea slug assemblage at fine spatial scales. By evaluating species richness, diversity indices, dominance patterns, and relationships with microhabitat conditions, we provide the first quantitative baseline for ongoing and future monitoring of sea slug communities in this warming region. This study establishes a framework for detecting temporal biodiversity shifts in temperate coastal ecosystems and offers new insights into the spatial–environmental drivers shaping benthic community dynamics under climate change.

## MATERIALS AND METHODS

### Study sites

Surveys were conducted at two coastal sites in northwestern Kyushu, Japan—Tatsunokuchi and Nomozaki–Akase, both located in Nagasaki Prefecture (Fig. 1 and Supplementary Fig. S1). These sites represent typical warm-temperate rocky reef habitats characterized by bedrock, boulders, and small sandy patches. The shallow subtidal zone supports diverse benthic invertebrate assemblages and well-developed macroalgal communities, providing suitable conditions for year-round observation of sea slugs.

**Fig. 1.**
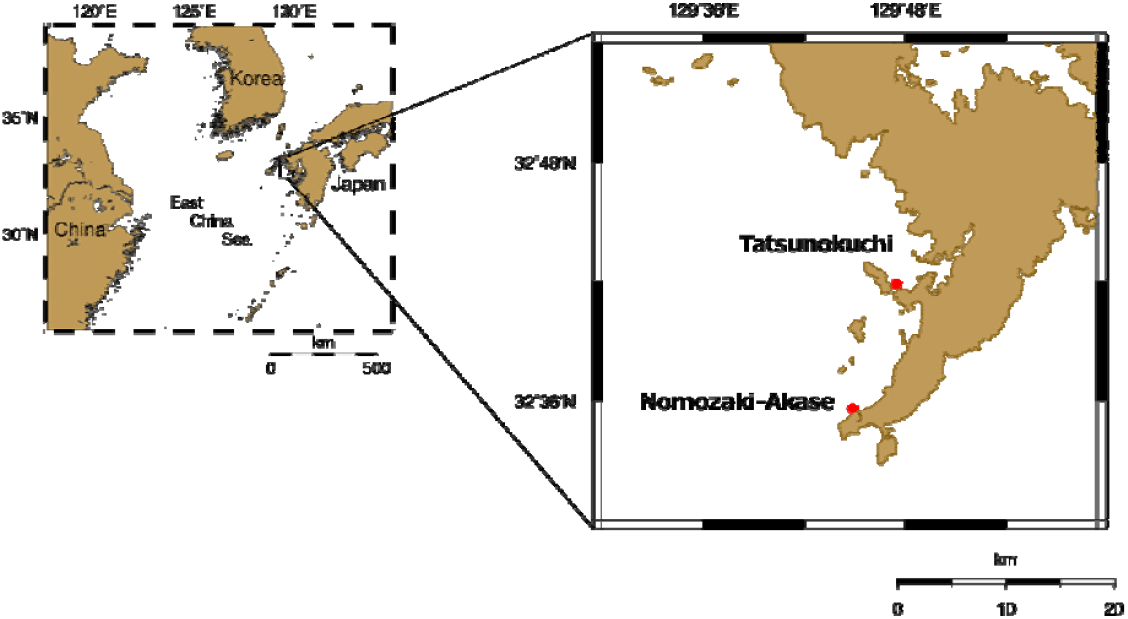
Survey sites of sea-slug (Heterobranchia) assemblages in the north west coast, Kyushu, Japan. Photographs of study sites (Tatsunokuchi and Nomozaki-Akase) are shown in Supplementary Fig. S1.

The transects spanned depths ranging from approximately 3 to 15 m (Fig. 2). Approximate depths at low tide were 10 m (Transect A), 8 m (Transect B), 3 m (Transect C), and 4 m (Transect D), with tidal variation of approximately ±1–2 m. Both sites were selected based on their ecological representativeness, accessibility for continuous SCUBA-based monitoring, and permission granted by local fisheries cooperatives.

**Fig. 2.**
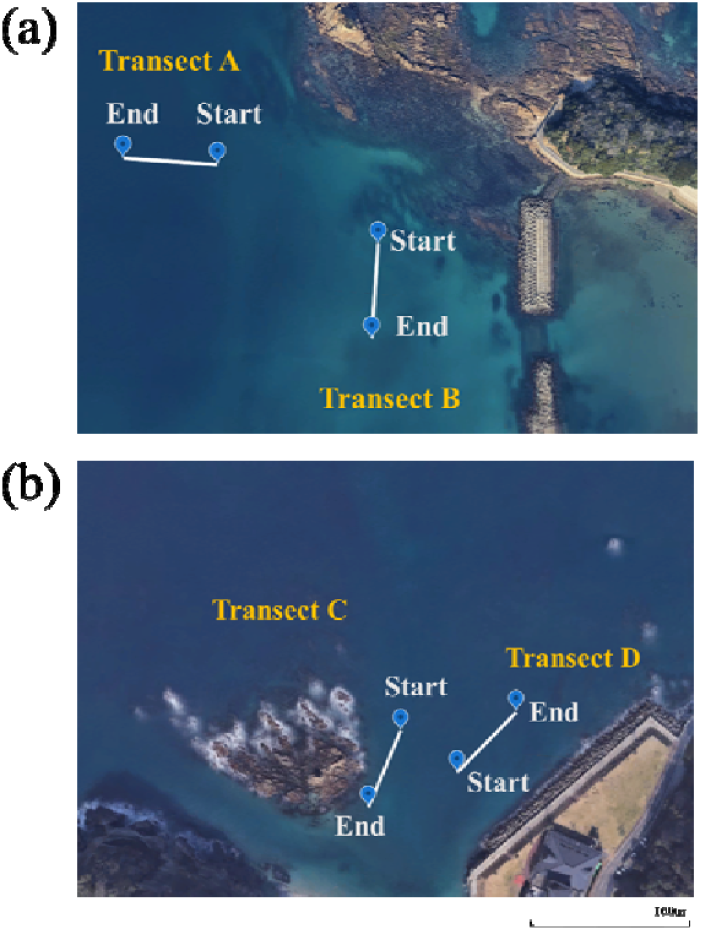
Locations of the line transects at (a) Tatsunokuchi and (b) Nomozaki–Akase in the north west coast, Kyushu, Japan. Base map and satellite imagery sourced from Google Earth (© Google, © 2025 Maxar Technologies).

### SCUBA diving survey

Monthly SCUBA surveys were conducted from February 2024 to November 2025 at the two study sites (Tatsunokuchi and Nomozaki–Akase) (Supplementary Fig. S2). Surveys were carried out across a wide range of tidal conditions, including both high and low tides. In total, 56 dives were completed, resulting in 61.25 hours of underwater observation. During each dive, in situ water temperature estimated using a diving computer (TUSA, IQ1204), underwater visibility, and general substrate characteristics were recorded immediately after surfacing (Supplementary Table S1). Tidal height and tidal cycle (spring, neap, and intermediate tides) were obtained from official tide tables corresponding to each survey date (Supplementary Table S1).

For quantitative sampling, four 50-m transects were established for repeated surveys: two at Tatsunokuchi (Transects A and B) and two at Nomozaki–Akase (Transects C and D) (Fig. 2). Transects were positioned to encompass multiple microhabitats, including rocky reefs, sandy patches, gravel substrates, and macroalgal beds. Because permanent underwater markers could not be installed due to fisheries and environmental regulations, transects were relocated during each survey using distinctive geomorphological features. The latitude and longitude of the start and end points of each transect are provided in Table 1. All sea slugs observed within 2 m on either side of the transect line (total belt width: 4 m) were recorded. Each survey was conducted by two divers, with one diver assigned to each side of the transect to avoid overlapping observations. Surveys at each site consisted of one or two dives, with a total bottom time of 40–80 minutes per sampling occasion.

Sea slugs were primarily documented by *in situ* underwater photography. Individuals whose diagnostic morphological features were clearly visible in photographs were identified solely based on images. In contrast, very small individuals, specimens photographed under poor underwater visibility, or individuals requiring confirmation of critical morphological characters for reliable identification were collected in the minimum number necessary and brought to the laboratory for examination under a stereomicroscope.Underwater photographs were taken using a digital camera (OLYMPUS Tough TG-6). Collected specimens were observed in detail under a stereomicroscope after each survey.

Species identification was based on morphological comparisons using the identification guides of Nakano (2019) and Gosliner et al. (2008). Taxonomic nomenclature and distributional records were verified using the World Register of Marine Species (WoRMS) and relevant primary literature, and supplemented by regional records reported by Ono and Kato (2020). Specimens that could not be confidently identified to species level were recorded as “cf. species name” or “sp.”

This study was conducted for academic research purposes under permits issued by the Nagasaki Prefectural Fisheries Division (Permit Nos. 6-Gyoshin-kyo 68 and 69; 7-Gyoshin-kyo 65 and 66). All diving surveys were conducted under the supervision of a certified diving instructor to ensure safety.

### Data analysis

Species diversity was quantified following Mittelbach and McGill (2023) using species richness, the Shannon–Wiener diversity index (H′), and the Simpson diversity index (D). Relative dominance was calculated as the proportion of individuals of each species within each sample. Because H′ is sensitive to both species evenness and overall diversity, whereas D emphasizes dominance and is less influenced by rare species, the combined use of these indices allowed a comprehensive assessment of temporal changes in heterobranch sea slug assemblage structure.

Relationships between water temperature and species richness or total abundance were examined separately for each study site using Spearman’s rank correlation coefficient (ρ) based on monthly observations. Water temperature values used in these analyses were recorded in situ during each diving survey using a diving computer. All correlation analyses were conducted as two-tailed tests.

All statistical analyses were conducted using R version 4.4.2 (R Core Team, 2024), with statistical significance evaluated at an alpha level of 0.05.

## RESULTS

During the study period from February 2024 to November 2025, a total of 81 sea slug species comprising 892 individuals were recorded. Representative photographs of nudibranch species recorded during the surveys are shown in Supplementary Fig. S3. At the site level, 47 species and 510 individuals were observed at Tatsunokuchi, whereas 56 species and 382 individuals were recorded at Nomozaki– Akase. Among the four transects, Transect A yielded 46 species and 411 individuals, Transect B 23 species and 97 individuals, Transect C 44 species and 239 individuals, and Transect D 38 species and 143 individuals. Detailed species- and transect-level data are provided in Supplementary Table S2.

At Tatsunokuchi (Fig. 3a), *Chromodoris orientalis* (20.5%) and *Hypselodoris festiva* (16.5%) were particularly dominant overall, followed by *Doriprismatica atromarginata, Dermatobranchus primus*, and *Goniobranchus sinensis*. The dominance of *C. orientali*s and *H. festiva* was especially pronounced in summer, whereas their relative contributions were lower in winter–spring when additional species were more evenly represented. At Nomozaki–Akase (Fig. 3b), *Aplysia japonica* (18.1%) exhibited the highest relative abundance, driven by the occurrence of large numbers of individuals on a limited number of survey dates, followed by *Dendrodoris krusensternii* (17.5%), which was also dominant. Similar to Tatsunokuchi, assemblages at Nomozaki–Akase exhibited a seasonal transition toward stronger dominance by a few species in summer, whereas winter–spring communities were more even. Full species-level and seasonal abundance data are provided in Supplementary Table S3.

**Fig. 3.**
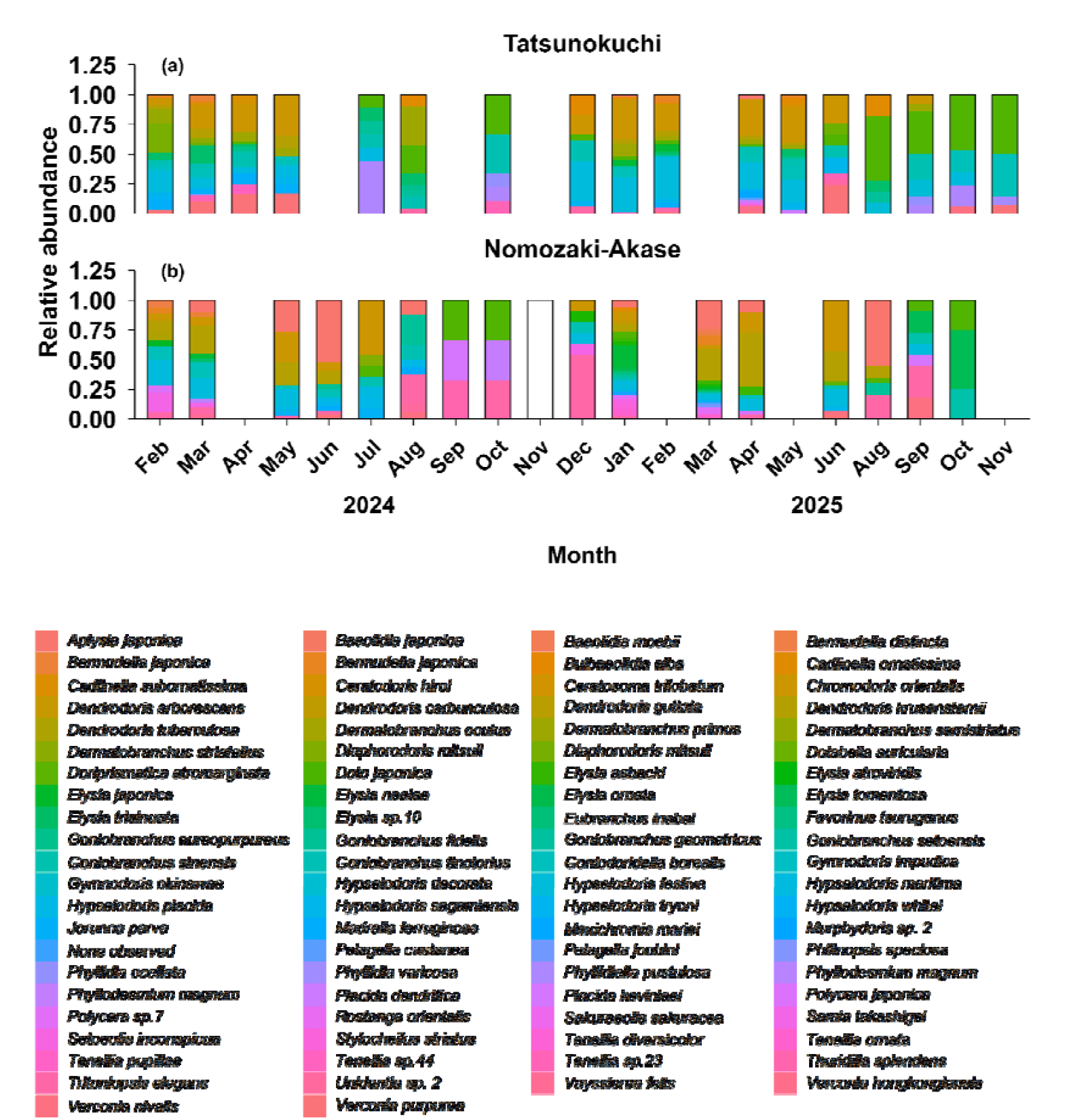
Seasonal variation in species composition of sea slug assemblages at (a) Tatsunokuchi and (b) Nomozaki–Akase, illustrating the shift from a more even community structure in winter– spring to dominance by a few species in summer.

Consistent with the observed seasonal shifts in species composition, species richness and diversity indices exhibited pronounced seasonal variation at both sites (Fig. 4). Species richness showed a concordant seasonal pattern. Nomozaki–Akase recorded its annual maximum of 24 species in January 2025, whereas Tatsunokuchi reached 18 species in February 2025 (Fig. 4a). Diversity was generally higher in winter–spring and declined markedly during summer. At Tatsunokuchi, the Shannon–Wiener index (H′) reached a maximum of 2.55 in March 2024, followed by a decline to 1.4–1.6 during summer (July–October) (Fig. 4b). A similar pattern was observed at Nomozaki–Akase, where H′ peaked at 2.49 in March 2024 and decreased to approximately 1.10 in autumn (Fig. 4b). The Simpson diversity index (D) varied in parallel with H′: at Tatsunokuchi, D was highest in spring (approximately 0.90) and declined to 0.64–0.74 in summer, while Nomozaki–Akase also exhibited its lowest D values during summer (Fig. 4c). These patterns indicate a seasonal transition from a more even assemblage structure in winter–spring to a dominance-driven structure in summer at both sites.

**Fig. 4.**
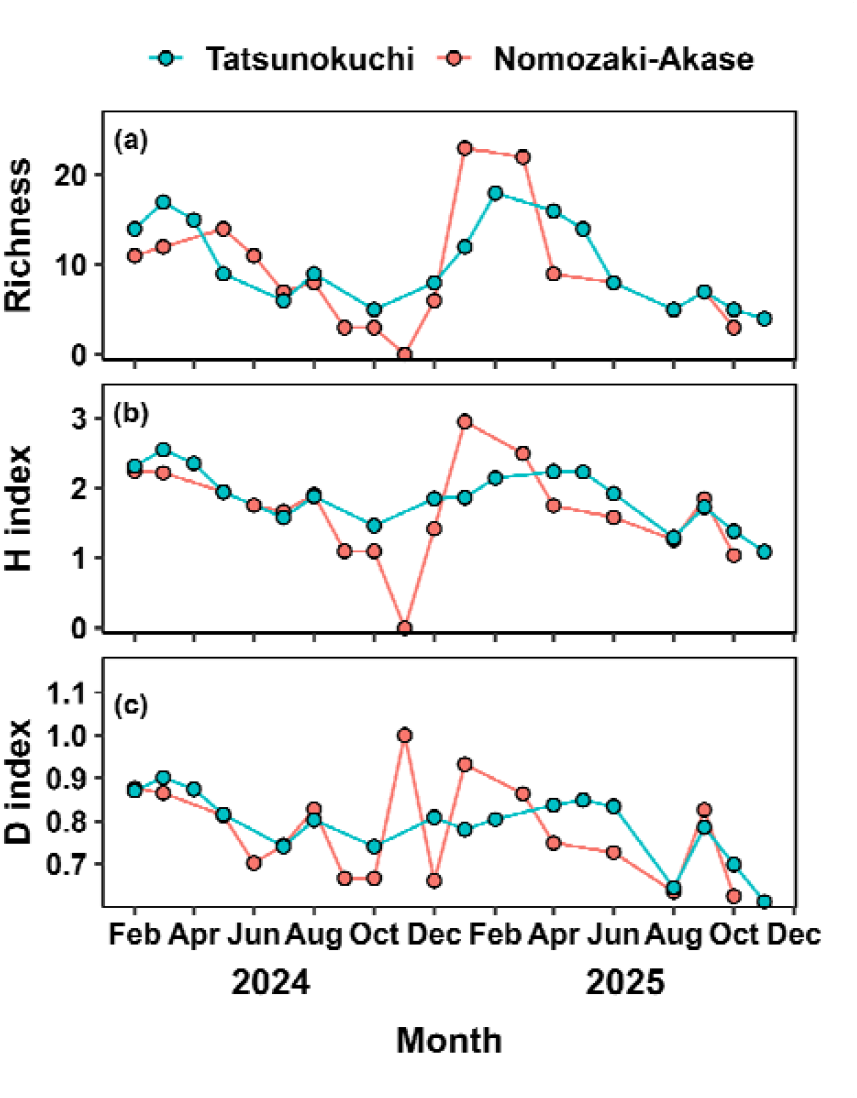
Monthly variation in species richness (a) and diversity indices (Shannon index (b) and Simpson index (c)) at each study site.

Water temperature exhibited clear seasonal variation at both sites (Fig. 5). Monthly mean temperatures derived from fixed temperature loggers declined to approximately 13–15 °C in winter (February–March), increased from spring through summer, and reached ∼28–29 °C in August–September, before decreasing again from autumn to early winter. The annual temperature range was approximately 14–15 °C at both sites. Nomozaki–Akase tended to experience slightly higher temperatures than Tatsunokuchi from summer to autumn, while differences among multiple loggers within the same site were small and seasonal patterns were generally consistent.

**Fig. 5.**
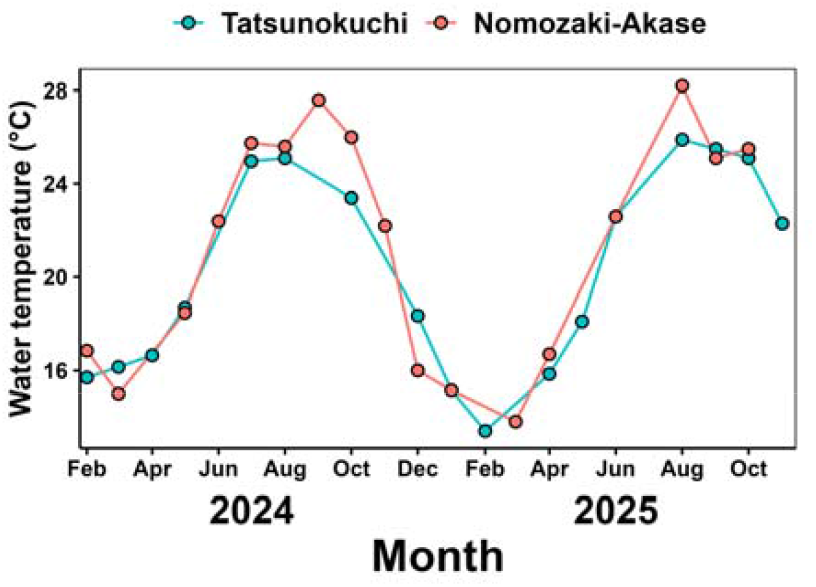
Monthly variation in water temperature recorded by diving computers from April 2024 to November 2025.

Water temperatures recorded during each diving survey were used to evaluate relationships with biological observations. Water temperature showed a significant negative relationship with both species richness and total abundance at both sites (Fig. 6). At Nomozaki–Akase, water temperature was negatively correlated with species richness (Spearman’s ρ = −0.708, p = 1.48 × 10^-3^) and total abundance (ρ = −0.629, p = 6.88 × 10^-3^). These relationships were even stronger at Tatsunokuchi, where increasing temperature was associated with pronounced declines in richness (ρ = −0.788, p = 1.72 × 10^-4^) and abundance (ρ = −0.854, p = 1.26 × 10^-5^). Together, these results indicate that seasonal temperature variation is closely associated with temporal changes in sea slug richness, abundance, and assemblage structure in the study area.

**Fig. 6.**
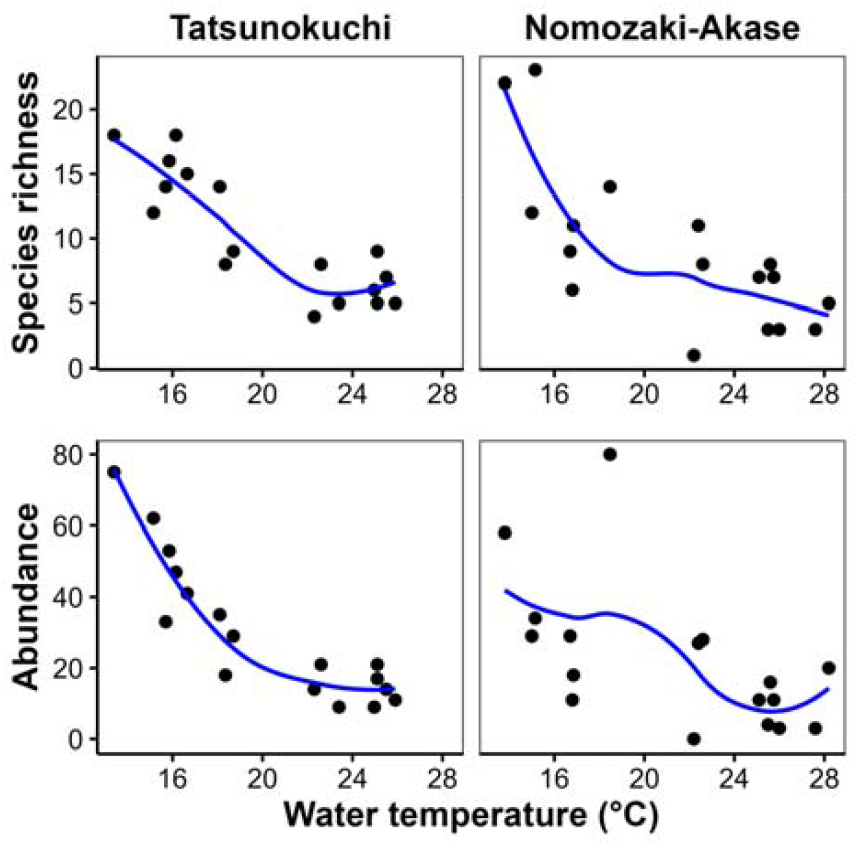
Relationships between water temperature and species richness and abundance of nudibranchs at the two study sites. Points represent monthly observations, and the blue line indicates a LOESS smoothing curve. Associations with water temperature were evaluated using Spearman’s rank correlation coefficient (ρ).

## DISCUSSION

In this study, nudibranch communities at two coastal sites in Nagasaki Prefecture, Japan exhibited site-specific differences in species composition and relative dominance, while showing a common seasonal response pattern to water temperature. Specifically, species richness, diversity, and abundance increased during winter to spring and declined markedly in summer at both sites. Pronounced seasonal dynamics are a common feature of benthic invertebrate communities in temperate coastal ecosystems, where environmental constraints fluctuate strongly over the annual cycle (Thompson, 1976; Todd, 1981).

Seasonal variation in water temperature is considered the primary driver of the observed community patterns. For marine ectotherms, temperature strongly constrains physiological performance, metabolic balance, and survival (Pörtner, 2002; Pörtner & Farrell, 2008). In temperate systems, summer water temperatures may approach or exceed the upper thermal tolerance of many species, resulting in reduced activity and increased mortality (Sunday et al., 2012). Consistent with these general patterns, our results revealed strong negative correlations between water temperature and both species richness and abundance, indicating that elevated summer temperatures represent a major limiting factor for nudibranch communities in this region. In contrast, the increase in diversity during winter to spring likely reflects convergence toward optimal thermal conditions for temperate species, combined with seasonal increases in food availability.

Diversity indices (Shannon and Simpson) exhibited clear seasonal variation similar to that observed for species richness, although the underlying mechanisms differed according to the properties of each index. During summer, overall abundance declined while a small number of dominant species (e.g., *Hypselodoris festiva* and *Hypselodoris crassicornis*) accounted for a relatively large proportion of individuals, resulting in reduced evenness and a pronounced decrease in the Shannon index. Changes in evenness and dominance structure are known to strongly influence diversity metrics and often accompany environmental stress in benthic communities (Magurran, 2004). The Simpson index, which is particularly sensitive to dominance, captured the seasonal transition from a relatively even community in winter to spring to a simplified community dominated by few species in summer.

Despite the shared seasonal response to water temperature, differences in relative dominance between sites suggest that additional factors beyond temperature contribute to community dynamics. At the Nomozaki-Akase site, green algae proliferated conspicuously during winter to spring, coinciding with an increase in sacoglossan nudibranchs. Sacoglossans are well known for their close trophic associations with specific green algal taxa, often exhibiting strong host specificity (Jensen, 1997; Christa et al., 2015; Morelli et al., 2023). The observed site-specific patterns therefore likely reflect differences in the seasonal availability of food resources and algal habitat structure. Such bottom-up effects of primary producers on consumer communities have been widely documented in benthic ecosystems (Underwood & Chapman, 1996), and our results indicate that these processes can modulate seasonal dynamics even within the same climatic region.

The seasonal pattern observed in this temperate system contrasts with reports from subtropical regions such as northern Taiwan, where nudibranch species richness increases during warmer periods (Chan et al., 2022). This contrast suggests that seasonal responses of nudibranch communities vary along latitudinal gradients, reflecting differences in thermal regimes and life-history constraints. Across marine ectotherms, thermal optima and tolerance limits often shift with latitude, resulting in region-specific responses to seasonal warming (Sunday et al., 2012; Deutsch et al., 2015). In temperate waters such as those off Nagasaki, Japan, elevated temperatures likely impose physiological stress, whereas in subtropical regions, higher temperatures may facilitate growth and recruitment. This apparent reversal highlights the strong dependence of nudibranch seasonal ecology on regional oceanographic and climatic conditions.

Several limitations of this study should be acknowledged. Environmental variables other than water temperature (e.g., current regimes, wave exposure, and seasonal variation in sessile prey organisms) were not explicitly evaluated, and thus community changes cannot be attributed solely to temperature effects. In addition, survey methods based on visual observations and photographic records may underestimate small or cryptic species, particularly those associated with encrusting substrates, and issues related to unidentified taxa or species complexes remain (Gosliner et al., 2018). Nevertheless, by providing multi-year, site-resolved, species-level data, this study offers a valuable foundation for long-term and broad-scale investigations of nudibranch community dynamics in temperate coastal ecosystems.

Importantly, the results of this study provide a quantitative description of the current seasonal structure of nudibranch communities in a temperate coastal region and represent an important baseline for evaluating future environmental change. Coastal ecosystems worldwide are experiencing rapid warming and increasing thermal variability (IPCC, 2023), and shifts in phenology, species composition, and dominance structure are expected to intensify. Detecting such changes requires well-documented reference data describing community states prior to major environmental alterations. By applying consistent survey methods across multiple sites and years, this study establishes a robust benchmark against which future changes in nudibranch community structure can be assessed. Such baseline data are particularly valuable given the scarcity of long-term, species-level observations for visually conspicuous but understudied benthic taxa such as nudibranchs.

## Acknowledgements

We are deeply grateful to Mr. Koji Sugisaki of the dive shop *Smilers* (https://smilers.jp/) for his dedicated assistance and for helping ensure safe and smooth field operations during our underwater surveys. We also thank the Nomozaki Sanwa Fisheries Cooperative and the Seihi Southern Fisheries Cooperative for kindly granting access to the survey sites and for their continued collaboration. We further appreciate the constructive feedback provided by the journal editors and anonymous reviewers, which significantly improved this study. This research was funded by the Sasakawa Scientific Research Grant from the Japan Science Society (Grant Number: 2024-4058).

## Ethics Approval

The research required no permit approvals.

## Author Contributions

RK Investigation, Writing – original draft, Visualization, Funding acquisition. MY Conceptualization, Methodology, Investigation, Writing – original draft, Supervision, Funding acquisition, Visualization.

## Supporting Information

Additional supporting information can be found online in the Supporting Information.

## Data Availability

Data has been provided as Supplementary Material.

## References

Chan, H.Y., Chang, Y.W., Chen, L.S., Nishida, K. & Shao, Y.T. 2024. The effect of environmental factors on spatial–temporal variation of heterobranch sea slug community in northern Taiwan. Frontiers in Marine Science, 11: 1374526. 10.3389/fmars.2024.1374526

Chou, H.N., Chen, W.C. & Chen, K.B. 2022. The effect of environmental factors on spatial–temporal variation of sea slug communities. Frontiers in Marine Science, 9: 1042961. 10.3389/fmars.2022.1042961

Christa, G., Händeler, K., Kloas, W. & Wägele, H. 2015. Acquired phototrophy through retention of functional chloroplasts increases growth efficiency of the sea slug Elysia viridis. PLOS ONE, 10: e0128729. 10.1371/journal.pone.0128729

Dinapoli, A. & Klussmann-Kolb, A. 2010. The long way to diversity—phylogeny and evolution of the Heterobranchia (Mollusca: Gastropoda). Molecular Phylogenetics and Evolution, 55: 60–76. 10.1016/j.ympev.2009.09.019

Deutsch, C., Ferrel, A., Seibel, B., Portner, H. O., & Takasuka, A. (2015). Climate change tightens a metabolic constraint on marine habitats. Science, 348(6239), 1132–1135. 10.1126/science.aaa1605

Gosliner, T.M., Behrens, D.W. & Valdés, Á. 2008. Indo-Pacific nudibranchs and sea slugs: a field guide to the world’s most diverse fauna. Monterey, CA: Sea Challengers Natural History Books & California Academy of Sciences.

Gosliner, T.M., Valdés, Á. & Behrens, D.W. 2018. Nudibranch and sea slug identification challenges in a changing ocean. Marine Biodiversity, 48: 1–14. 10.1007/s12526-018-0890-1

Hongo, C. & Yamano, H. 2013. Species-specific responses of corals to bleaching in a high-latitude region (Tatsukushi, Shikoku Island, Japan). Palaeogeography, Palaeoclimatology, Palaeoecology, 392: 274– 284.

IPCC. 2023. Climate Change 2023: Synthesis Report. Cambridge University Press.

Jensen, K.R. 1997. Evolution of the Sacoglossa (Mollusca, Opisthobranchia) and the ecological associations with their food plants. Evolutionary Ecology, 11: 301–335.

Kashio, S., Kawase, M., Ukai, H., Ooya, M., Nishi, H. & Asada, K. 2021. Intertidal nudibranchs in Minamichita Town, Aichi Prefecture I (Nudibranchia). Nagoya Biodiversity, 8: 1–22.

Kato, R. & Yagi, M. 2025. A different world: temporal changes in nudibranch community structure over a half-century. PeerJ. (in press).

Kumagai, N.H., Yamakita, T., Watanabe, K., Tanaka, K., Hidaka, M., Kakinuma, Y. & Yamano, H. 2018. Ocean currents and herbivory drive macroalgae-to-coral community shift under climate warming. Proceedings of the National Academy of Sciences of the United States of America, 115: 8606–8611. 10.1073/pnas.1716826115

Laetz, E.M.J., et al. 2024. Critical thermal maxima and oxygen uptake in Elysia viridis, a sea slug that steals chloroplasts to photosynthesize. Journal of Experimental Biology, 227: jeb246331. 10.1242/jeb.246331

Magurran, A.E. 2004. Measuring biological diversity. Oxford: Blackwell Publishing.

Mittelbach, G.G. & McGill, B.J. 2023. Community ecology, 2nd edn. Tokyo: Maruzen Publishing.

Morelli, L., Vitale, L., Mollo, E., Polese, G. & Mazzella, V. 2023. Food-shaped photosynthesis: photophysiology of the sea slug Elysia viridis fed with two alternative chloroplast donors. Marine Biology, 170: 140.

Nakano, R. 2019. Sea slugs of Japan, 2nd edn. Tokyo: Bun-ichi Sogo Shuppan. (in Japanese)

Nimbs, A.L. & Smith, S.D.A. 2015. Range extensions for heterobranch sea slugs on the eastern coast of Australia. Marine Biodiversity Records, 8: e122. 10.1017/S1755267215000524

Nimbs, M.J. & Smith, S.D.A. 2018. Beyond Capricornia: tropical sea slugs extend their distributions into the Tasman Sea. Diversity, 10: 99. 10.3390/d10030099

Ono, A. & Kato, S. 2020. Illustrated guide to 1260 species of sea slugs. Tokyo: Seibundo Shinkosha. (in Japanese)

Ota, Y., Tamura, S., Yamasaki, E., Togawa, Y. & Nakano, R. 2021. Sea slugs from the eastern coast of Tottori Prefecture and surrounding waters. Bulletin of the Tottori Prefectural Museum, 58: 1–47.

Pinsky, M.L., Selden, R.L. & Kitchel, Z.J. 2020. Climate-driven shifts in marine species ranges: scaling from organisms to communities. Annual Review of Marine Science, 12: 153–179. 10.1146/annurev-marine-010419-010916

Poloczanska, E.S., Brown, C.J., Sydeman, W.J., Kiessling, W., Schoeman, D.S., Moore, P.J., Brander, K.M., Bruno, J.F., Buckley, L.B. & Richardson, A.J. 2013. Global imprint of climate change on marine life. Nature Climate Change, 3: 919–925. 10.1038/nclimate1958

Pörtner, H. O. 2002. Physiological basis of temperature-dependent biogeography: trade-offs in metabolic adaptation and relevance for species’ sensitivity to climate change. Journal of Experimental Biology, 205(15), 2217–2230. 10.1242/jeb.205.15.2217

Pörtner, H. O., & Farrell, A. P. 2008. Physiology and climate change. Science, 322(5902), 690–692.10.1126/science.1163156

R Core Team. 2024. R:A Language and Environment for Statistical Computing. R Foundation for Statistical Computing, Vienna, Austria. URL https://www.R-project.org/.

Rees, H.L., Eggleton, J.D., Rachor, E. & Vanden Berghe, E. 2007. Structure and dynamics of the North Sea benthos. ICES Cooperative Research Report No. 288.

Sunday, J.M., Bates, A.E. & Dulvy, N.K. 2012. Thermal tolerance and the global redistribution of animals. Nature Climate Change, 2: 686–690.

Thompson, R. J. (1976). The influence of diet and low seston concentration on energy budget and reproductive effort of Mytilus edulis. Marine Biology, 35(1), 83–93. 10.1371/journal.pone.0109796

Todd, C. D. (1981). The ecology of nudibranch molluscs. Oceanography and Marine Biology: An Annual Review, 19, 141–234.

Underwood, A. J., & Chapman, M. G. (1996). Scales of spatial patterns of distribution of intertidal invertebrates. Oecologia, 107(2), 212–224. 10.1007/BF00327905

Vergés, A., Steinberg, P.D., Hay, M.E., Poore, A.G.B., Campbell, A.H., Ballesteros, E. & Wilson, S.K. 2014. The tropicalization of temperate marine ecosystems. Proceedings of the Royal Society B, 281: 20140846. 10.1098/rspb.2014.0846

Wernberg, T., Bennett, S., Babcock, R.C., de Bettignies, T., Cure, K., Depczynski, M., Dufois, F., Fromont, J., Fulton, C.J., Hovey, R.K., Harvey, E.S., Holmes, T.H., Kendrick, G.A., Radford, B., Santana-Garcon, J., Saunders, B.J., Smale, D.A., Thomsen, M.S., Tuckett, C.A., Tuya, F., Vanderklift, M.A., Wilson, S., 2016. Climate-driven regime shift of a temperate marine ecosystem. Science (80-.). 353, 169–172. 10.1126/science.aad8745

WoRMS Editorial Board. 2026. World Register of Marine Species. Available at https://www.marinespecies.org (accessed 1 January 2026). 10.14284/170

Yasuhara, M., Doi, H., Wei, C.-L., Danovaro, R., Myhre, S.E., Tittensor, D.P. & Rogers, A.D. 2020. Past and future decline of tropical pelagic biodiversity. Proceedings of the National Academy of Sciences of the United States of America, 117: 12891–12896. 10.1073/pnas.1916923117

