## Supplementary Information for "Quantitative assessment of sea slugs (Heterobranchia) assemblages along the western coast of Kyushu, Japan: a baseline for long-term biodiversity monitoring"

**Table S1** Summary of environmental data recorded during the present scuba diving surveys.

| **Date** | **Site** | **Line** | **Duration_min** | **Temp_air_c** | **Temp_water_c** | **Depth_avg_m** | **Depth_max_m** | **Transparency_m** | **Wave_state** | **Tide** | **Weather** |
| --- | --- | --- | --- | --- | --- | --- | --- | --- | --- | --- | --- |
| 2024-02-06 | Tatsunokuchi | B | 69 | 7.3 | 15.3 | 4.5 | 6.3 | 8 | Moderate tide (ebb phase) | with swell | Cloudy |
| 2024-02-06 | Tatsunokuchi | A | 58 | 12.6 | 16.1 | 11.3 | 15.5 | 8 | Moderate tide (low tide) | with swell | Sunny |
| 2024-03-28 | Tatsunokuchi | B | 59 | 12.7 | 16.2 | 12.3 | 17 | 6 | Moderate tide (high tide) | with swell | Rainy |
| 2024-03-28 | Tatsunokuchi | A | 39 | 18 | 16.1 | 4.8 | 7.8 | 6 | Moderate tide (high tide) | with swell | Rainy |
| 2024-04-18 | Tatsunokuchi | B | 67 | 17.9 | 16.3 | 11.2 | 14.1 | 3 | Neap tide (low tide) | no swell | Sunny |
| 2024-04-18 | Tatsunokuchi | A | 52 | 21.8 | 17 | 5.4 | 6.5 | 3 | Neap tide (low tide) | no swell | Sunny |
| 2024-05-24 | Tatsunokuchi | B | 60 | 23.4 | 18.2 | 11.3 | 15.2 | 4-6 | Spring tide (high tide) | no swell | Cloudy |
| 2024-05-24 | Tatsunokuchi | A | 44 | 24.4 | 19.2 | 5 | 6.7 | 4-6 | Spring tide (ebb phase) | no swell | Cloudy |
| 2024-07-10 | Tatsunokuchi | B | 42 | 22.6 | 25.8 | 5.6 | 7.6 | 4-6 | Moderate tide | with swell | Cloudy |
| 2024-07-10 | Tatsunokuchi | A | 58 | 27.2 | 24.2 | 10.8 | 15 | 4-6 | Moderate tide | with swell | Cloudy |
| 2024-07-26 | Tatsunokuchi | A・B | 82 | 30.2 | 24.9 | 9.5 | 15.4 | 8 | Moderate tide | with swell | Sunny |
| 2024-08-26 | Tatsunokuchi | A・B | 92 | 27.7 | 25.1 | 8.7 | 14.8 | 3 | Neap tide | with swell | Sunny |
| 2024-10-25 | Tatsunokuchi | A・B | 89 | 21.1 | 23.4 |  |  | 4-6 | Moderate tide (low tide) | with swell | Cloudy |
| 2024-12-09 | Tatsunokuchi | B | 43 | 9 | 18.3 |  | 15.2 | 10 | Neap tide (low tide) | no swell | Cloudy |
| 2024-12-09 | Tatsunokuchi | A | 40 | 9 | 18.4 |  | 6.4 | 10 | Neap tide (low tide) | no swell | Cloudy |
| 2025-01-27 | Tatsunokuchi | B | 58 | 8 | 15.4 | 10.9 | 14.6 | 8 | Moderate tide (ebb phase) | no swell | Rainy |
| 2025-01-27 | Tatsunokuchi | A | 43 | 9.7 | 14.9 | 5.4 | 7.1 | 8 | Moderate tide (ebb phase) | no swell | Rainy |
| 2025-02-28 | Tatsunokuchi | B | 70 | 17 | 13.3 | 11.7 | 16.4 | 8 | Spring tide (high tide) | with swell | Cloudy |
| 2025-02-28 | Tatsunokuchi | A | 47 | 18 | 13.5 | 5.4 | 7 | 8 | Spring tide (high tide) | no swell | Cloudy |
| 2025-04-11 | Tatsunokuchi | B | 74 | 21 | 15.8 | 10.6 | 15.8 | 3 | Spring tide (ebb phase) | no swell | Sunny |
| 2025-04-11 | Tatsunokuchi | A | 58 | 21 | 15.9 | 4.7 | 6.5 | 3 | Spring tide (ebb phase) | no swell | Sunny |
| 2025-05-26 | Tatsunokuchi | B | 52 | 23 | 17.7 | 10.6 | 15.3 | 4-6 | Spring tide (ebb phase) | with current | Cloudy |
| 2025-05-26 | Tatsunokuchi | A | 54 | 24 | 18.5 | 4.6 | 6.9 | 4-6 | Spring tide (ebb phase) | with current | Cloudy |
| 2025-06-27 | Tatsunokuchi | A・B | 86 | 31 | 22.6 | 8.9 | 16.6 | 3 | Moderate tide (high tide) | with current | Cloudy |
| 2025-08-25 | Tatsunokuchi | A・B | 97 | 33 | 25.9 | 9.3 | 15.3 | 8 | Moderate tide (high tide) | no swell | Sunny |
| 2025-09-29 | Tatsunokuchi | A・B | 87 | 28 | 25.5 | 10.1 | 21.3 | 6-8 | Neap tide (ebb phase) | with current | Cloudy |
| 2025-10-17 | Tatsunokuchi | A・B | 84 | 30 | 25.1 | 9.3 | 19.2 | 6-8 | Young tide (ebb phase) | with current | Sunny |
| 2025-11-04 | Tatsunokuchi | A・B | 82 | 19 | 22.3 | 9.3 | 16.1 | 4-6 | Spring tide (ebb phase) | with current | Sunny |
| 2024-02-05 | Nomozaki-Akase | D | 65 | 11.9 | 16.7 | 4 | 6.1 | 5-7 | Young tide (low tide) | with swell | Rainy |
| 2024-02-05 | Nomozaki-Akase | C | 39 | 14.1 | 17 | 3.9 | 6 | 5-7 | Young tide (low tide) | with swell | Cloudy |
| 2024-03-27 | Nomozaki-Akase | D | 54 | 12 | 14.8 | 4.4 | 7.1 | 4-6 | Moderate tide (high tide) | with swell | Sunny |
| 2024-03-27 | Nomozaki-Akase | C | 54 | 15.4 | 15.2 | 3.5 | 4.5 | 4-6 | Moderate tide (ebb phase) | with swell | Sunny |
| 2024-05-08 | Nomozaki-Akase | D | 66 | 17.5 | 18.3 | 3.9 | 6.5 | 6-8 | Moderate tide (ebb phase) | with swell | Cloudy |
| 2024-05-08 | Nomozaki-Akase | C | 40 | 18.1 | 18.2 | 2.8 | 3.7 | 6-8 | Moderate tide (low tide) | with swell | Cloudy |
| 2024-05-23 | Nomozaki-Akase | D | 67 | 20.4 | 18.6 | 3.5 | 6.1 | 8 | Spring tide (high tide) | with swell | Cloudy |
| 2024-05-23 | Nomozaki-Akase | C | 44 | 22.2 | 18.8 | 2.9 | 3.9 | 8 | Spring tide (ebb phase) | with swell | Rainy |
| 2024-06-17 | Nomozaki-Akase | D | 65 | 24.2 | 22.3 | 3.3 | 5.7 | 4 | Moderate tide | with swell | Cloudy |
| 2024-06-17 | Nomozaki-Akase | C | 43 | 26.5 | 22.5 | 3.3 | 4.3 | 8 | Moderate tide | with swell | Cloudy |
| 2024-07-23 | Nomozaki-Akase | D | 73 | 30.1 | 25.5 | 4.6 | 7.3 | 6-8 | Moderate tide (high tide) | no swell | Sunny |
| 2024-07-23 | Nomozaki-Akase | C | 49 | 32.9 | 26 | 3.8 | 5.2 | 6-8 | Moderate tide (high tide) | no swell | Sunny |
| 2024-08-06 | Nomozaki-Akase | C・D | 106 | 31.9 | 25.6 | 4.2 | 7.2 | 8 | Moderate tide (high tide) | no swell | Sunny |
| 2024-09-11 | Nomozaki-Akase | C・D | 92 | 29.7 | 27.6 | 3.6 | 6.1 | 5-7 | Neap tide (low tide) | with swell | Cloudy |
| 2024-10-07 | Nomozaki-Akase | C・D | 82 | 22.5 | 26 |  |  | 5-7 | Moderate tide (flood phase) | with swell | Cloudy |
| 2024-11-12 | Nomozaki-Akase | C・D | 78 | 22.9 | 22.2 | 2.9 | 5.2 | 6 | Moderate tide (flood phase) | no swell | Sunny |
| 2025-12-20 | Nomozaki-Akase | D | 59 | 6 | 15.2 | 4.2 | 6.6 | 3 | Moderate tide (flood phase) | with swell | Cloudy |
| 2024-12-20 | Nomozaki-Akase | C | 40 | 12.9 | 16.8 | 4.1 | 5.3 | 3 | Moderate tide (high tide) | with swell | Cloudy |
| 2025-01-14 | Nomozaki-Akase | D | 68 | 11 | 15.7 | 4.3 |  | 4-6 | Spring tide (high tide) | with swell | Cloudy |
| 2025-01-14 | Nomozaki-Akase | C | 63 | 12 | 14.6 | 3.7 | 4.8 | 4-6 | Spring tide (high tide) | with swell | Cloudy |
| 2025-03-10 | Nomozaki-Akase | D | 75 | 14 | 13.8 | 4 | 6.3 | 6-8 | Young tide | with swell | Cloudy |
| 2025-03-10 | Nomozaki-Akase | C | 72 | 15 | 13.8 | 3.6 | 4.4 | 6-8 | Young tide | with swell | Rainy |
| 2025-04-10 | Nomozaki-Akase | D | 70 | 21 | 16.1 | 3.6 | 6.3 | 4-6 | Moderate tide | with swell | Rainy |
| 2025-04-10 | Nomozaki-Akase | C | 59 | 21 | 17.3 | 3.3 | 4.2 | 4-6 | Moderate tide | with swell | Rainy |
| 2025-06-26 | Nomozaki-Akase | C・D | 95 | 29 | 22.6 | 4 | 7 | 5 | Spring tide (high tide) | with swell | Cloudy |
| 2025-08-18 | Nomozaki-Akase | C・D | 84 | 33 | 28.2 |  | 5.4 | 5 | Long tide (low tide) | with swell | Rainy |
| 2025-09-22 | Nomozaki-Akase | C・D | 94 | 28 | 25.1 | 4.4 | 7.2 | 3 | Spring tide (high tide) | strong swell | Cloudy |
| 2025-10-10 | Nomozaki-Akase | C・D | 93 | 28.7 | 25.5 | 4.9 | 7.6 | 6 | Moderate tide (high tide) | with swell | Sunny |

**Table S2** Summary of heterobranch sea-slug assemblages recorded during the present scuba diving surveys.

| **Date** | **Site** | **Line** | **Species** | **Count** |
| --- | --- | --- | --- | --- |
| 2024/2/6 | Tatsunokuchi | A | *Diaphorodoris mitsuii* | 8 |
| 2024/2/6 | Tatsunokuchi | A | *Dermatobranchus primus* | 4 |
| 2024/2/6 | Tatsunokuchi | A | *Goniobranchus sinensis* | 1 |
| 2024/2/6 | Tatsunokuchi | A | *Goniobranchus tinctorius* | 1 |
| 2024/2/6 | Tatsunokuchi | A | *Gymnodoris impudica* | 1 |
| 2024/2/6 | Tatsunokuchi | A | *Verconia purpurea* | 1 |
| 2024/2/6 | Tatsunokuchi | A | *Jorunna parva* | 2 |
| 2024/2/6 | Tatsunokuchi | A | *Dendrodoris krusensternii* | 1 |
| 2024/2/6 | Tatsunokuchi | A | *Hypselodoris festiva* | 3 |
| 2024/2/6 | Tatsunokuchi | A | *Bermudella japonica* | 1 |
| 2024/2/6 | Tatsunokuchi | B | *Hypselodoris sagamiensis* | 1 |
| 2024/2/6 | Tatsunokuchi | B | *Hypselodoris festiva* | 3 |
| 2024/2/6 | Tatsunokuchi | B | *Jorunna parva* | 1 |
| 2024/2/6 | Tatsunokuchi | B | *Elysia trisinuata* | 2 |
| 2024/2/6 | Tatsunokuchi | B | *Chromodoris orientalis* | 2 |
| 2024/2/6 | Tatsunokuchi | B | *Hypselodoris tryoni* | 1 |
| 2024/3/28 | Tatsunokuchi | A | *Verconia purpurea* | 5 |
| 2024/3/28 | Tatsunokuchi | A | *Hypselodoris festiva* | 2 |
| 2024/3/28 | Tatsunokuchi | A | Dendrodoris guttata | 1 |
| 2024/3/28 | Tatsunokuchi | A | *Ceratodoris hiroi* | 1 |
| 2024/3/28 | Tatsunokuchi | A | *Goniobranchus sinensis* | 2 |
| 2024/3/28 | Tatsunokuchi | A | *Bermudella japonica* | 2 |
| 2024/3/28 | Tatsunokuchi | A | *Chromodoris orientalis* | 7 |
| 2024/3/28 | Tatsunokuchi | A | *Hypselodoris tryoni* | 1 |
| 2024/3/28 | Tatsunokuchi | A | *Dermatobranchus semistriatus* | 2 |
| 2024/3/28 | Tatsunokuchi | A | *Goniobranchus tinctorius* | 3 |
| 2024/3/28 | Tatsunokuchi | A | *Dendrodoris krusensternii* | 1 |
| 2024/3/28 | Tatsunokuchi | A | *Tenellia* sp.23 | 2 |
| 2024/3/28 | Tatsunokuchi | A | *Goniodoridella borealis* | 1 |
| 2024/3/28 | Tatsunokuchi | A | *Diaphorodoris mitsuii* | 1 |
| 2024/3/28 | Tatsunokuchi | B | *Bermudella japonica* | 1 |
| 2024/3/28 | Tatsunokuchi | B | *Jorunna parva* | 1 |
| 2024/3/28 | Tatsunokuchi | B | *Chromodoris orientalis* | 2 |
| 2024/3/28 | Tatsunokuchi | B | *Elysia trisinuata* | 7 |
| 2024/3/28 | Tatsunokuchi | B | *Dendrodoris krusensternii* | 2 |
| 2024/3/28 | Tatsunokuchi | B | *Hypselodoris festiva* | 2 |
| 2024/3/28 | Tatsunokuchi | B | *Philinopsis speciosa* | 1 |
| 2024/4/18 | Tatsunokuchi | A | *Goniobranchus sinensis* | 4 |
| 2024/4/18 | Tatsunokuchi | A | *Tenellia* sp.44 | 3 |
| 2024/4/18 | Tatsunokuchi | A | *Verconia purpurea* | 7 |
| 2024/4/18 | Tatsunokuchi | A | *Dermatobranchus primus* | 3 |
| 2024/4/18 | Tatsunokuchi | A | *Goniobranchus fidelis* | 1 |
| 2024/4/18 | Tatsunokuchi | A | *Chromodoris orientalis* | 9 |
| 2024/4/18 | Tatsunokuchi | A | *Hypselodoris maritima* | 1 |
| 2024/4/18 | Tatsunokuchi | A | *Favorinus tsuruganus* | 1 |
| 2024/4/18 | Tatsunokuchi | A | *Goniobranchus tinctorius* | 1 |
| 2024/4/18 | Tatsunokuchi | A | *Jorunna parva* | 3 |
| 2024/4/18 | Tatsunokuchi | A | *Cadlinella subornatissima* | 1 |
| 2024/4/18 | Tatsunokuchi | B | *Hypselodoris festiva* | 1 |
| 2024/4/18 | Tatsunokuchi | B | *Ceratodoris hiroi* | 2 |
| 2024/4/18 | Tatsunokuchi | B | *Jorunna parva* | 1 |
| 2024/4/18 | Tatsunokuchi | B | *Doriprismatica atromarginata* | 1 |
| 2024/4/18 | Tatsunokuchi | B | *Chromodoris orientalis* | 1 |
| 2024/4/18 | Tatsunokuchi | B | *Elysia trisinuata* | 1 |
| 2024/5/24 | Tatsunokuchi | A | *Jorunna parva* | 2 |
| 2024/5/24 | Tatsunokuchi | A | *Chromodoris orientalis* | 8 |
| 2024/5/24 | Tatsunokuchi | A | *Verconia purpurea* | 5 |
| 2024/5/24 | Tatsunokuchi | A | *Dermatobranchus primus* | 2 |
| 2024/5/24 | Tatsunokuchi | A | *Hypselodoris maritima* | 1 |
| 2024/5/24 | Tatsunokuchi | A | *Goniobranchus tinctorius* | 1 |
| 2024/5/24 | Tatsunokuchi | A | *Hypselodoris festiva* | 1 |
| 2024/5/24 | Tatsunokuchi | A | *Hypselodoris sagamiensis* | 1 |
| 2024/5/24 | Tatsunokuchi | B | *Dendrodoris krusensternii* | 3 |
| 2024/5/24 | Tatsunokuchi | B | *Chromodoris orientalis* | 2 |
| 2024/5/24 | Tatsunokuchi | B | *Hypselodoris sagamiensis* | 1 |
| 2024/5/24 | Tatsunokuchi | B | *Hypselodoris maritima* | 1 |
| 2024/5/24 | Tatsunokuchi | B | *Goniobranchus tinctorius* | 1 |
| 2024/7/10 | Tatsunokuchi | A | *Goniobranchus sinensis* | 1 |
| 2024/7/10 | Tatsunokuchi | A | *Phyllidiella pustulosa* | 1 |
| 2024/7/10 | Tatsunokuchi | B | *Phyllidiella pustulosa* | 2 |
| 2024/7/10 | Tatsunokuchi | B | *Doriprismatica atromarginata* | 1 |
| 2024/7/10 | Tatsunokuchi | B | *Goniobranchus fidelis* | 1 |
| 2024/7/26 | Tatsunokuchi | A | *Elysia trisinuata* | 1 |
| 2024/7/26 | Tatsunokuchi | A | *Phyllidiella pustulosa* | 1 |
| 2024/7/26 | Tatsunokuchi | A | *Hypselodoris maritima* | 1 |
| 2024/8/26 | Tatsunokuchi | A | *Cadlinella ornatissima* | 1 |
| 2024/8/26 | Tatsunokuchi | A | *Tritoniopsis elegans* | 1 |
| 2024/8/26 | Tatsunokuchi | A | *Dermatobranchus primus* | 7 |
| 2024/8/26 | Tatsunokuchi | A | *Doriprismatica atromarginata* | 3 |
| 2024/8/26 | Tatsunokuchi | A | *Goniobranchus sinensis* | 2 |
| 2024/8/26 | Tatsunokuchi | A | *Bulbaeolidia alba* | 1 |
| 2024/8/26 | Tatsunokuchi | B | *Goniobranchus aureopurpureus* | 1 |
| 2024/8/26 | Tatsunokuchi | B | *Doriprismatica atromarginata* | 2 |
| 2024/8/26 | Tatsunokuchi | B | *Elysia trisinuata* | 2 |
| 2024/8/26 | Tatsunokuchi | B | *Goniobranchus fidelis* | 1 |
| 2024/10/25 | Tatsunokuchi | A | *Goniobranchus sinensis* | 3 |
| 2024/10/25 | Tatsunokuchi | A | *Doriprismatica atromarginata* | 3 |
| 2024/10/25 | Tatsunokuchi | A | *Phyllidiella pustulosa* | 1 |
| 2024/10/25 | Tatsunokuchi | A | *Phyllidia ocellata* | 1 |
| 2024/10/25 | Tatsunokuchi | A | *Tenellia* sp.23 | 1 |
| 2024/12/9 | Tatsunokuchi | A | *Goniobranchus sinensis* | 2 |
| 2024/12/9 | Tatsunokuchi | A | *Tritoniopsis elegans* | 1 |
| 2024/12/9 | Tatsunokuchi | A | *Hypselodoris festiva* | 5 |
| 2024/12/9 | Tatsunokuchi | A | *Chromodoris orientalis* | 2 |
| 2024/12/9 | Tatsunokuchi | A | *Bulbaeolidia alba* | 1 |
| 2024/12/9 | Tatsunokuchi | B | *Cadlinella ornatissima* | 2 |
| 2024/12/9 | Tatsunokuchi | B | *Hypselodoris festiva* | 1 |
| 2024/12/9 | Tatsunokuchi | B | *Chromodoris orientalis* | 1 |
| 2024/12/9 | Tatsunokuchi | B | *Goniobranchus sinensis* | 1 |
| 2024/12/9 | Tatsunokuchi | B | *Doriprismatica atromarginata* | 1 |
| 2024/12/9 | Tatsunokuchi | B | *Hypselodoris tryoni* | 1 |
| 2025/1/27 | Tatsunokuchi | A | *Hypselodoris festiva* | 17 |
| 2025/1/27 | Tatsunokuchi | A | *Dermatobranchus primus* | 6 |
| 2025/1/27 | Tatsunokuchi | A | *Elysia japonica* | 1 |
| 2025/1/27 | Tatsunokuchi | A | *Goniobranchus tinctorius* | 4 |
| 2025/1/27 | Tatsunokuchi | A | *Goniobranchus sinensis* | 1 |
| 2025/1/27 | Tatsunokuchi | A | *Chromodoris orientalis* | 18 |
| 2025/1/27 | Tatsunokuchi | A | *Dendrodoris krusensternii* | 1 |
| 2025/1/27 | Tatsunokuchi | A | *Tenellia* sp.23 | 1 |
| 2025/1/27 | Tatsunokuchi | A | *Doriprismatica atromarginata* | 1 |
| 2025/1/27 | Tatsunokuchi | A | *Goniobranchus geometricus* | 1 |
| 2025/1/27 | Tatsunokuchi | A | *Aplysia japonica* | 1 |
| 2025/1/27 | Tatsunokuchi | B | *Chromodoris orientalis* | 3 |
| 2025/1/27 | Tatsunokuchi | B | *Dendrodoris krusensternii* | 2 |
| 2025/1/27 | Tatsunokuchi | B | *Elysia japonica* | 2 |
| 2025/1/27 | Tatsunokuchi | B | *Cadlinella ornatissima* | 1 |
| 2025/1/27 | Tatsunokuchi | B | *Hypselodoris festiva* | 1 |
| 2025/1/27 | Tatsunokuchi | B | *Doriprismatica atromarginata* | 1 |
| 2025/2/28 | Tatsunokuchi | A | *Goniobranchus aureopurpureus* | 1 |
| 2025/2/28 | Tatsunokuchi | A | *Hypselodoris festiva* | 23 |
| 2025/2/28 | Tatsunokuchi | A | *Chromodoris orientalis* | 17 |
| 2025/2/28 | Tatsunokuchi | A | *Dermatobranchus primus* | 2 |
| 2025/2/28 | Tatsunokuchi | A | *Dermatobranchus oculus* | 1 |
| 2025/2/28 | Tatsunokuchi | A | *Doriprismatica atromarginata* | 2 |
| 2025/2/28 | Tatsunokuchi | A | *Hypselodoris sagamiensis* | 2 |
| 2025/2/28 | Tatsunokuchi | A | *Verconia purpurea* | 1 |
| 2025/2/28 | Tatsunokuchi | A | *Eubranchus inabai* | 1 |
| 2025/2/28 | Tatsunokuchi | A | *Elysia atroviridis* | 1 |
| 2025/2/28 | Tatsunokuchi | A | *Tenellia* sp.23 | 1 |
| 2025/2/28 | Tatsunokuchi | A | *Dendrodoris krusensternii* | 2 |
| 2025/2/28 | Tatsunokuchi | A | *Jorunna parva* | 3 |
| 2025/2/28 | Tatsunokuchi | A | *Bermudella japonica* | 5 |
| 2025/2/28 | Tatsunokuchi | A | *Tritoniopsis elegans* | 2 |
| 2025/2/28 | Tatsunokuchi | B | Elysia tomentosa | 1 |
| 2025/2/28 | Tatsunokuchi | B | *Hypselodoris festiva* | 4 |
| 2025/2/28 | Tatsunokuchi | B | *Elysia japonica* | 4 |
| 2025/2/28 | Tatsunokuchi | B | *Dendrodoris krusensternii* | 1 |
| 2025/2/28 | Tatsunokuchi | B | *Cadlinella ornatissima* | 1 |
| 2025/4/11 | Tatsunokuchi | A | *Verconia purpurea* | 1 |
| 2025/4/11 | Tatsunokuchi | A | *Dendrodoris krusensternii* | 2 |
| 2025/4/11 | Tatsunokuchi | A | *Chromodoris orientalis* | 13 |
| 2025/4/11 | Tatsunokuchi | A | *Aplysia japonica* | 1 |
| 2025/4/11 | Tatsunokuchi | A | *Goniobranchus sinensis* | 4 |
| 2025/4/11 | Tatsunokuchi | A | *Doriprismatica atromarginata* | 1 |
| 2025/4/11 | Tatsunokuchi | A | *Mexichromis mariei* | 3 |
| 2025/4/11 | Tatsunokuchi | A | *Hypselodoris festiva* | 10 |
| 2025/4/11 | Tatsunokuchi | A | *Pelagella castanea* | 1 |
| 2025/4/11 | Tatsunokuchi | A | *Dermatobranchus primus* | 2 |
| 2025/4/11 | Tatsunokuchi | A | *Rostanga orientalis* | 1 |
| 2025/4/11 | Tatsunokuchi | A | *Jorunna parva* | 1 |
| 2025/4/11 | Tatsunokuchi | A | *Tenellia* sp.23 | 1 |
| 2025/4/11 | Tatsunokuchi | A | *Goniobranchus tinctorius* | 3 |
| 2025/4/11 | Tatsunokuchi | A | *Verconia hongkongiensis* | 2 |
| 2025/4/11 | Tatsunokuchi | B | *Aplysia japonica* | 1 |
| 2025/4/11 | Tatsunokuchi | B | *Chromodoris orientalis* | 3 |
| 2025/4/11 | Tatsunokuchi | B | *Hypselodoris festiva* | 2 |
| 2025/4/11 | Tatsunokuchi | B | *Placida dendritica* | 1 |
| 2025/5/26 | Tatsunokuchi | A | *Hypselodoris festiva* | 6 |
| 2025/5/26 | Tatsunokuchi | A | *Chromodoris orientalis* | 8 |
| 2025/5/26 | Tatsunokuchi | A | *Goniobranchus tinctorius* | 4 |
| 2025/5/26 | Tatsunokuchi | A | *Elysia trisinuata* | 2 |
| 2025/5/26 | Tatsunokuchi | A | *Goniobranchus sinensis* | 2 |
| 2025/5/26 | Tatsunokuchi | A | *Cadlinella ornatissima* | 1 |
| 2025/5/26 | Tatsunokuchi | A | *Ceratosoma trilobatum* | 1 |
| 2025/5/26 | Tatsunokuchi | A | *Bermudella japonica* | 1 |
| 2025/5/26 | Tatsunokuchi | A | *Dermatobranchus primus* | 1 |
| 2025/5/26 | Tatsunokuchi | A | *Bulbaeolidia alba* | 1 |
| 2025/5/26 | Tatsunokuchi | A | *Hypselodoris sagamiensis* | 2 |
| 2025/5/26 | Tatsunokuchi | A | *Goniobranchus aureopurpureus* | 1 |
| 2025/5/26 | Tatsunokuchi | B | *Dendrodoris krusensternii* | 1 |
| 2025/5/26 | Tatsunokuchi | B | *Placida dendritica* | 1 |
| 2025/5/26 | Tatsunokuchi | B | *Hypselodoris festiva* | 1 |
| 2025/5/26 | Tatsunokuchi | B | *Chromodoris orientalis* | 2 |
| 2025/6/27 | Tatsunokuchi | A | *Doriprismatica atromarginata* | 2 |
| 2025/6/27 | Tatsunokuchi | A | *Chromodoris orientalis* | 5 |
| 2025/6/27 | Tatsunokuchi | A | *Hypselodoris sagamiensis* | 3 |
| 2025/6/27 | Tatsunokuchi | A | *Verconia purpurea* | 5 |
| 2025/6/27 | Tatsunokuchi | A | *Goniobranchus sinensis* | 2 |
| 2025/6/27 | Tatsunokuchi | A | *Diaphorodoris mitsuii* | 2 |
| 2025/6/27 | Tatsunokuchi | A | *Tritoniopsis elegans* | 1 |
| 2025/6/27 | Tatsunokuchi | A | *Tenellia* sp.23 | 1 |
| 2025/8/25 | Tatsunokuchi | A | *Elysia trisinuata* | 1 |
| 2025/8/25 | Tatsunokuchi | A | *Cadlinella ornatissima* | 2 |
| 2025/8/25 | Tatsunokuchi | A | *Doriprismatica atromarginata* | 5 |
| 2025/8/25 | Tatsunokuchi | A | *Hypselodoris decorata* | 1 |
| 2025/8/25 | Tatsunokuchi | A | *Goniobranchus fidelis* | 1 |
| 2025/8/25 | Tatsunokuchi | B | *Doriprismatica atromarginata* | 1 |
| 2025/9/29 | Tatsunokuchi | A | *Dermatobranchus primus* | 1 |
| 2025/9/29 | Tatsunokuchi | A | *Doriprismatica atromarginata* | 4 |
| 2025/9/29 | Tatsunokuchi | A | *Goniobranchus sinensis* | 3 |
| 2025/9/29 | Tatsunokuchi | A | *Phyllidiella pustulosa* | 1 |
| 2025/9/29 | Tatsunokuchi | A | *Hypselodoris decorata* | 2 |
| 2025/9/29 | Tatsunokuchi | A | *Phyllidia ocellata* | 1 |
| 2025/9/29 | Tatsunokuchi | A | *Chromodoris orientalis* | 1 |
| 2025/9/29 | Tatsunokuchi | B | *Doriprismatica atromarginata* | 1 |
| 2025/10/17 | Tatsunokuchi | A | *Verconia nivalis* | 1 |
| 2025/10/17 | Tatsunokuchi | A | *Doriprismatica atromarginata* | 7 |
| 2025/10/17 | Tatsunokuchi | A | *Goniobranchus tinctorius* | 3 |
| 2025/10/17 | Tatsunokuchi | A | *Hypselodoris festiva* | 2 |
| 2025/10/17 | Tatsunokuchi | A | *Phyllidiella pustulosa* | 2 |
| 2025/10/17 | Tatsunokuchi | B | *Doriprismatica atromarginata* | 1 |
| 2025/10/17 | Tatsunokuchi | B | *Phyllidiella pustulosa* | 1 |
| 2025/11/4 | Tatsunokuchi | A | *Doriprismatica atromarginata* | 5 |
| 2025/11/4 | Tatsunokuchi | A | *Goniobranchus tinctorius* | 5 |
| 2025/11/4 | Tatsunokuchi | A | *Phyllidia varicosa* | 1 |
| 2025/11/4 | Tatsunokuchi | A | *Verconia purpurea* | 1 |
| 2025/11/4 | Tatsunokuchi | B | *Doriprismatica atromarginata* | 2 |
| 2024/2/5 | Nomozaki-Akase | C | *Elysia atroviridis* | 1 |
| 2024/2/5 | Nomozaki-Akase | C | *Hypselodoris festiva* | 2 |
| 2024/2/5 | Nomozaki-Akase | C | *Dendrodoris krusensternii* | 1 |
| 2024/2/5 | Nomozaki-Akase | C | *Goniobranchus tinctorius* | 2 |
| 2024/2/5 | Nomozaki-Akase | C | *Chromodoris orientalis* | 1 |
| 2024/2/5 | Nomozaki-Akase | C | *Polycera japonica* | 1 |
| 2024/2/5 | Nomozaki-Akase | C | *Samla takashigei* | 2 |
| 2024/2/5 | Nomozaki-Akase | C | *Bermudella distincta* | 1 |
| 2024/2/5 | Nomozaki-Akase | C | *Bermudella japonica* | 1 |
| 2024/2/5 | Nomozaki-Akase | D | *Tritoniopsis elegans* | 1 |
| 2024/2/5 | Nomozaki-Akase | D | *Hypselodoris festiva* | 2 |
| 2024/2/5 | Nomozaki-Akase | D | *Dendrodoris krusensternii* | 1 |
| 2024/2/5 | Nomozaki-Akase | D | *Dendrodoris tuberculosa* | 1 |
| 2024/2/5 | Nomozaki-Akase | D | *Samla takashigei* | 1 |
| 2024/3/27 | Nomozaki-Akase | C | *Tritoniopsis elegans* | 1 |
| 2024/3/27 | Nomozaki-Akase | C | *Goniobranchus tinctorius* | 2 |
| 2024/3/27 | Nomozaki-Akase | C | *Dendrodoris krusensternii* | 1 |
| 2024/3/27 | Nomozaki-Akase | C | *Hypselodoris festiva* | 3 |
| 2024/3/27 | Nomozaki-Akase | C | *Samla takashigei* | 1 |
| 2024/3/27 | Nomozaki-Akase | C | *Elysia japonica* | 1 |
| 2024/3/27 | Nomozaki-Akase | C | *Phyllodesmium magnum* | 1 |
| 2024/3/27 | Nomozaki-Akase | C | *Bermudella japonica* | 1 |
| 2024/3/27 | Nomozaki-Akase | D | *Chromodoris orientalis* | 1 |
| 2024/3/27 | Nomozaki-Akase | D | *Goniobranchus tinctorius* | 2 |
| 2024/3/27 | Nomozaki-Akase | D | *Aplysia japonica* | 3 |
| 2024/3/27 | Nomozaki-Akase | D | *Tritoniopsis elegans* | 1 |
| 2024/3/27 | Nomozaki-Akase | D | *Dendrodoris krusensternii* | 6 |
| 2024/3/27 | Nomozaki-Akase | D | *Elysia trisinuata* | 1 |
| 2024/3/27 | Nomozaki-Akase | D | *Chromodoris orientalis* | 1 |
| 2024/3/27 | Nomozaki-Akase | D | *Vayssierea felis* | 1 |
| 2024/3/27 | Nomozaki-Akase | D | *Hypselodoris festiva* | 2 |
| 2024/5/8 | Nomozaki-Akase | C | *Hypselodoris festiva* | 2 |
| 2024/5/8 | Nomozaki-Akase | C | *Dendrodoris krusensternii* | 3 |
| 2024/5/8 | Nomozaki-Akase | C | *Chromodoris orientalis* | 8 |
| 2024/5/8 | Nomozaki-Akase | C | *Aplysia japonica* | 3 |
| 2024/5/8 | Nomozaki-Akase | C | *Goniobranchus tinctorius* | 1 |
| 2024/5/8 | Nomozaki-Akase | D | *Chromodoris orientalis* | 3 |
| 2024/5/8 | Nomozaki-Akase | D | *Dendrodoris krusensternii* | 2 |
| 2024/5/8 | Nomozaki-Akase | D | *Hypselodoris sagamiensis* | 1 |
| 2024/5/8 | Nomozaki-Akase | D | *Madrella ferruginosa* | 1 |
| 2024/5/8 | Nomozaki-Akase | D | *Aplysia japonica* | 1 |
| 2024/5/8 | Nomozaki-Akase | D | *Hypselodoris festiva* | 1 |
| 2024/5/8 | Nomozaki-Akase | D | *Goniobranchus sinensis* | 1 |
| 2024/5/23 | Nomozaki-Akase | C | *Tritoniopsis elegans* | 1 |
| 2024/5/23 | Nomozaki-Akase | C | *Hypselodoris festiva* | 6 |
| 2024/5/23 | Nomozaki-Akase | C | *Aplysia japonica* | 16 |
| 2024/5/23 | Nomozaki-Akase | C | *Verconia purpurea* | 1 |
| 2024/5/23 | Nomozaki-Akase | C | *Ceratodoris hiroi* | 1 |
| 2024/5/23 | Nomozaki-Akase | C | *Chromodoris orientalis* | 6 |
| 2024/5/23 | Nomozaki-Akase | C | *Hypselodoris placida* | 1 |
| 2024/5/23 | Nomozaki-Akase | C | *Dendrodoris krusensternii* | 6 |
| 2024/5/23 | Nomozaki-Akase | D | *Hypselodoris festiva* | 2 |
| 2024/5/23 | Nomozaki-Akase | D | *Dendrodoris krusensternii* | 5 |
| 2024/5/23 | Nomozaki-Akase | D | *Aplysia japonica* | 1 |
| 2024/5/23 | Nomozaki-Akase | D | *Goniobranchus tinctorius* | 1 |
| 2024/5/23 | Nomozaki-Akase | D | *Chromodoris orientalis* | 2 |
| 2024/5/23 | Nomozaki-Akase | D | *Jorunna parva* | 1 |
| 2024/5/23 | Nomozaki-Akase | D | *Hypselodoris placida* | 1 |
| 2024/5/23 | Nomozaki-Akase | D | *Hypselodoris sagamiensis* | 1 |
| 2024/5/23 | Nomozaki-Akase | D | *Hypselodoris maritima* | 1 |
| 2024/6/17 | Nomozaki-Akase | C | *Aplysia japonica* | 13 |
| 2024/6/17 | Nomozaki-Akase | C | *Chromodoris orientalis* | 1 |
| 2024/6/17 | Nomozaki-Akase | C | *Dendrodoris krusensternii* | 3 |
| 2024/6/17 | Nomozaki-Akase | C | *Madrella ferruginosa* | 1 |
| 2024/6/17 | Nomozaki-Akase | C | *Goniobranchus sinensis* | 1 |
| 2024/6/17 | Nomozaki-Akase | C | *Verconia purpurea* | 1 |
| 2024/6/17 | Nomozaki-Akase | C | *Goniobranchus aureopurpureus* | 1 |
| 2024/6/17 | Nomozaki-Akase | C | *Hypselodoris sagamiensis* | 1 |
| 2024/6/17 | Nomozaki-Akase | D | *Hypselodoris maritima* | 1 |
| 2024/6/17 | Nomozaki-Akase | D | *Goniobranchus tinctorius* | 1 |
| 2024/6/17 | Nomozaki-Akase | D | *Aplysia japonica* | 1 |
| 2024/6/17 | Nomozaki-Akase | D | *Chromodoris orientalis* | 1 |
| 2024/6/17 | Nomozaki-Akase | D | *Tritoniopsis elegans* | 1 |
| 2024/7/23 | Nomozaki-Akase | C | *Chromodoris orientalis* | 4 |
| 2024/7/23 | Nomozaki-Akase | C | *Goniobranchus sinensis* | 1 |
| 2024/7/23 | Nomozaki-Akase | C | *Hypselodoris maritima* | 1 |
| 2024/7/23 | Nomozaki-Akase | C | *Hypselodoris whitei* | 1 |
| 2024/7/23 | Nomozaki-Akase | C | *Hypselodoris placida* | 1 |
| 2024/7/23 | Nomozaki-Akase | C | *Dermatobranchus striatellus* | 1 |
| 2024/7/23 | Nomozaki-Akase | D | *Doriprismatica atromarginata* | 1 |
| 2024/7/23 | Nomozaki-Akase | D | *Chromodoris orientalis* | 1 |
| 2024/8/6 | Nomozaki-Akase | C | *Goniobranchus fidelis* | 4 |
| 2024/8/6 | Nomozaki-Akase | C | *Unidentia* sp. 2 | 1 |
| 2024/8/6 | Nomozaki-Akase | C | *Goniobranchus sinensis* | 1 |
| 2024/8/6 | Nomozaki-Akase | C | *Madrella ferruginosa* | 1 |
| 2024/8/6 | Nomozaki-Akase | C | *Tritoniopsis elegans* | 1 |
| 2024/8/6 | Nomozaki-Akase | D | *Tritoniopsis elegans* | 3 |
| 2024/8/6 | Nomozaki-Akase | D | *Hypselodoris placida* | 1 |
| 2024/8/6 | Nomozaki-Akase | D | *Aplysia japonica* | 2 |
| 2024/8/6 | Nomozaki-Akase | D | *Verconia nivalis* | 1 |
| 2024/8/6 | Nomozaki-Akase | D | *Goniobranchus sinensis* | 1 |
| 2024/9/11 | Nomozaki-Akase | C | *Placida kevinleei* | 1 |
| 2024/9/11 | Nomozaki-Akase | D | *Doriprismatica atromarginata* | 1 |
| 2024/9/11 | Nomozaki-Akase | D | *Thuridilla splendens* | 1 |
| 2024/10/7 | Nomozaki-Akase | C | *Tritoniopsis elegans* | 1 |
| 2024/10/7 | Nomozaki-Akase | C | *Phyllodesmium magnum* | 1 |
| 2024/10/7 | Nomozaki-Akase | C | *Doriprismatica atromarginata* | 1 |
| 2024/11/12 | Nomozaki-Akase | C・D | None observed | 0 |
| 2024/12/20 | Nomozaki-Akase | C | *Elysia asbecki* | 1 |
| 2024/12/20 | Nomozaki-Akase | C | *Chromodoris orientalis* | 1 |
| 2024/12/20 | Nomozaki-Akase | C | *Samla takashigei* | 1 |
| 2024/12/20 | Nomozaki-Akase | C | *Tritoniopsis elegans* | 2 |
| 2024/12/20 | Nomozaki-Akase | D | *Tritoniopsis elegans* | 4 |
| 2024/12/20 | Nomozaki-Akase | D | *Hypselodoris festiva* | 1 |
| 2024/12/20 | Nomozaki-Akase | D | *Goniobranchus sinensis* | 1 |
| 2025/1/14 | Nomozaki-Akase | C | *Elysia japonica* | 4 |
| 2025/1/14 | Nomozaki-Akase | C | *Polycera japonica* | 1 |
| 2025/1/14 | Nomozaki-Akase | C | *Doriprismatica atromarginata* | 1 |
| 2025/1/14 | Nomozaki-Akase | C | *Tenellia ornata* | 1 |
| 2025/1/14 | Nomozaki-Akase | C | *Stylocheilus striatus* | 1 |
| 2025/1/14 | Nomozaki-Akase | C | *Doto japonica* | 1 |
| 2025/1/14 | Nomozaki-Akase | C | *Tenellia diversicolor* | 1 |
| 2025/1/14 | Nomozaki-Akase | C | *Murphydoris* sp. 2 | 1 |
| 2025/1/14 | Nomozaki-Akase | C | *Chromodoris orientalis* | 2 |
| 2025/1/14 | Nomozaki-Akase | C | *Bulbaeolidia alba* | 1 |
| 2025/1/14 | Nomozaki-Akase | C | *Samla takashigei* | 1 |
| 2025/1/14 | Nomozaki-Akase | C | *Dendrodoris krusensternii* | 1 |
| 2025/1/14 | Nomozaki-Akase | C | *Hypselodoris maritima* | 1 |
| 2025/1/14 | Nomozaki-Akase | C | *Elysia* sp.10 | 1 |
| 2025/1/14 | Nomozaki-Akase | D | *Goniobranchus fidelis* | 1 |
| 2025/1/14 | Nomozaki-Akase | D | *Aplysia japonica* | 2 |
| 2025/1/14 | Nomozaki-Akase | D | *Tritoniopsis elegans* | 1 |
| 2025/1/14 | Nomozaki-Akase | D | *Dendrodoris krusensternii* | 1 |
| 2025/1/14 | Nomozaki-Akase | D | *Doriprismatica atromarginata* | 1 |
| 2025/1/14 | Nomozaki-Akase | D | *Setoeolis inconspicua* | 1 |
| 2025/1/14 | Nomozaki-Akase | D | *Hypselodoris festiva* | 2 |
| 2025/1/14 | Nomozaki-Akase | D | *Chromodoris orientalis* | 1 |
| 2025/1/14 | Nomozaki-Akase | D | *Elysia japonica* | 2 |
| 2025/1/14 | Nomozaki-Akase | D | *Elysia asbecki* | 1 |
| 2025/1/14 | Nomozaki-Akase | D | *Elysia nealae* | 1 |
| 2025/1/14 | Nomozaki-Akase | D | *Goniobranchus sinensis* | 1 |
| 2025/1/14 | Nomozaki-Akase | D | *Dendrodoris carbunculosa* | 1 |
| 2025/3/10 | Nomozaki-Akase | C | *Aplysia japonica* | 6 |
| 2025/3/10 | Nomozaki-Akase | C | *Dendrodoris krusensternii* | 13 |
| 2025/3/10 | Nomozaki-Akase | C | Gymnodoris okinawae | 1 |
| 2025/3/10 | Nomozaki-Akase | C | *Tenellia ornata* | 1 |
| 2025/3/10 | Nomozaki-Akase | C | *Bermudella japonica* | 5 |
| 2025/3/10 | Nomozaki-Akase | C | *Chromodoris orientalis* | 1 |
| 2025/3/10 | Nomozaki-Akase | C | *Sakuraeolis sakuracea* | 1 |
| 2025/3/10 | Nomozaki-Akase | C | *Bermudella distincta* | 2 |
| 2025/3/10 | Nomozaki-Akase | C | *Hypselodoris festiva* | 2 |
| 2025/3/10 | Nomozaki-Akase | C | *Polycera* sp.7 | 1 |
| 2025/3/10 | Nomozaki-Akase | C | *Elysia asbecki* | 1 |
| 2025/3/10 | Nomozaki-Akase | C | *Elysia atroviridis* | 1 |
| 2025/3/10 | Nomozaki-Akase | D | *Doto japonica* | 1 |
| 2025/3/10 | Nomozaki-Akase | D | *Jorunna parva* | 1 |
| 2025/3/10 | Nomozaki-Akase | D | *Aplysia japonica* | 7 |
| 2025/3/10 | Nomozaki-Akase | D | *Elysia* sp.10 | 1 |
| 2025/3/10 | Nomozaki-Akase | D | *Baeolidia japonica* | 1 |
| 2025/3/10 | Nomozaki-Akase | D | *Pelagella castanea* | 1 |
| 2025/3/10 | Nomozaki-Akase | D | *Doto japonica* | 1 |
| 2025/3/10 | Nomozaki-Akase | D | *Polycera* sp.7 | 1 |
| 2025/3/10 | Nomozaki-Akase | D | *Dendrodoris krusensternii* | 2 |
| 2025/3/10 | Nomozaki-Akase | D | *Chromodoris orientalis* | 1 |
| 2025/3/10 | Nomozaki-Akase | D | *Hypselodoris festiva* | 1 |
| 2025/3/10 | Nomozaki-Akase | D | *Polycera japonica* | 1 |
| 2025/3/10 | Nomozaki-Akase | D | *Tenellia pupillae* | 1 |
| 2025/3/10 | Nomozaki-Akase | D | *Pelagella joubini* | 1 |
| 2025/3/10 | Nomozaki-Akase | D | *Elysia japonica* | 1 |
| 2025/3/10 | Nomozaki-Akase | D | *Baeolidia moebii* | 1 |
| 2025/4/10 | Nomozaki-Akase | C | *Dendrodoris krusensternii* | 12 |
| 2025/4/10 | Nomozaki-Akase | C | *Tritoniopsis elegans* | 1 |
| 2025/4/10 | Nomozaki-Akase | C | *Chromodoris orientalis* | 3 |
| 2025/4/10 | Nomozaki-Akase | C | *Doto japonica* | 2 |
| 2025/4/10 | Nomozaki-Akase | C | *Hypselodoris festiva* | 2 |
| 2025/4/10 | Nomozaki-Akase | C | *Aplysia japonica* | 3 |
| 2025/4/10 | Nomozaki-Akase | C | *Goniobranchus tinctorius* | 1 |
| 2025/4/10 | Nomozaki-Akase | D | *Polycera japonica* | 1 |
| 2025/4/10 | Nomozaki-Akase | D | *Dendrodoris arborescens* | 1 |
| 2025/4/10 | Nomozaki-Akase | D | *Chromodoris orientalis* | 1 |
| 2025/4/10 | Nomozaki-Akase | D | *Hypselodoris festiva* | 1 |
| 2025/4/10 | Nomozaki-Akase | D | *Dendrodoris krusensternii* | 1 |
| 2025/6/26 | Nomozaki-Akase | C | *Dolabella auricularia* | 1 |
| 2025/6/26 | Nomozaki-Akase | C | *Hypselodoris festiva* | 2 |
| 2025/6/26 | Nomozaki-Akase | C | *Chromodoris orientalis* | 5 |
| 2025/6/26 | Nomozaki-Akase | C | *Goniobranchus sinensis* | 1 |
| 2025/6/26 | Nomozaki-Akase | C | *Verconia purpurea* | 1 |
| 2025/6/26 | Nomozaki-Akase | C | *Dendrodoris krusensternii* | 7 |
| 2025/6/26 | Nomozaki-Akase | D | *Hypselodoris festiva* | 2 |
| 2025/6/26 | Nomozaki-Akase | D | *Chromodoris orientalis* | 7 |
| 2025/6/26 | Nomozaki-Akase | D | *Hypselodoris sagamiensis* | 1 |
| 2025/6/26 | Nomozaki-Akase | D | *Verconia nivalis* | 1 |
| 2025/8/18 | Nomozaki-Akase | C | *Aplysia japonica* | 7 |
| 2025/8/18 | Nomozaki-Akase | C | *Goniobranchus fidelis* | 1 |
| 2025/8/18 | Nomozaki-Akase | C | *Dendrodoris krusensternii* | 1 |
| 2025/8/18 | Nomozaki-Akase | C | *Tritoniopsis elegans* | 2 |
| 2025/8/18 | Nomozaki-Akase | D | *Aplysia japonica* | 4 |
| 2025/8/18 | Nomozaki-Akase | D | *Doriprismatica atromarginata* | 1 |
| 2025/8/18 | Nomozaki-Akase | D | *Goniobranchus fidelis* | 1 |
| 2025/8/18 | Nomozaki-Akase | D | *Dendrodoris krusensternii* | 1 |
| 2025/8/18 | Nomozaki-Akase | D | *Tritoniopsis elegans* | 2 |
| 2025/9/22 | Nomozaki-Akase | C | *Verconia nivalis* | 1 |
| 2025/9/22 | Nomozaki-Akase | C | *Tritoniopsis elegans* | 3 |
| 2025/9/22 | Nomozaki-Akase | C | *Goniobranchus setoensis* | 1 |
| 2025/9/22 | Nomozaki-Akase | D | *Hypselodoris festiva* | 1 |
| 2025/9/22 | Nomozaki-Akase | D | *Verconia nivalis* | 1 |
| 2025/9/22 | Nomozaki-Akase | D | *Placida kevinleei* | 1 |
| 2025/9/22 | Nomozaki-Akase | D | *Elysia ornata* | 2 |
| 2025/9/22 | Nomozaki-Akase | D | *Doriprismatica atromarginata* | 1 |
| 2025/10/10 | Nomozaki-Akase | C | *Goniobranchus setoensis* | 1 |
| 2025/10/10 | Nomozaki-Akase | C | *Elysia ornata* | 2 |
| 2025/10/10 | Nomozaki-Akase | D | *Doriprismatica atromarginata* | 1 |

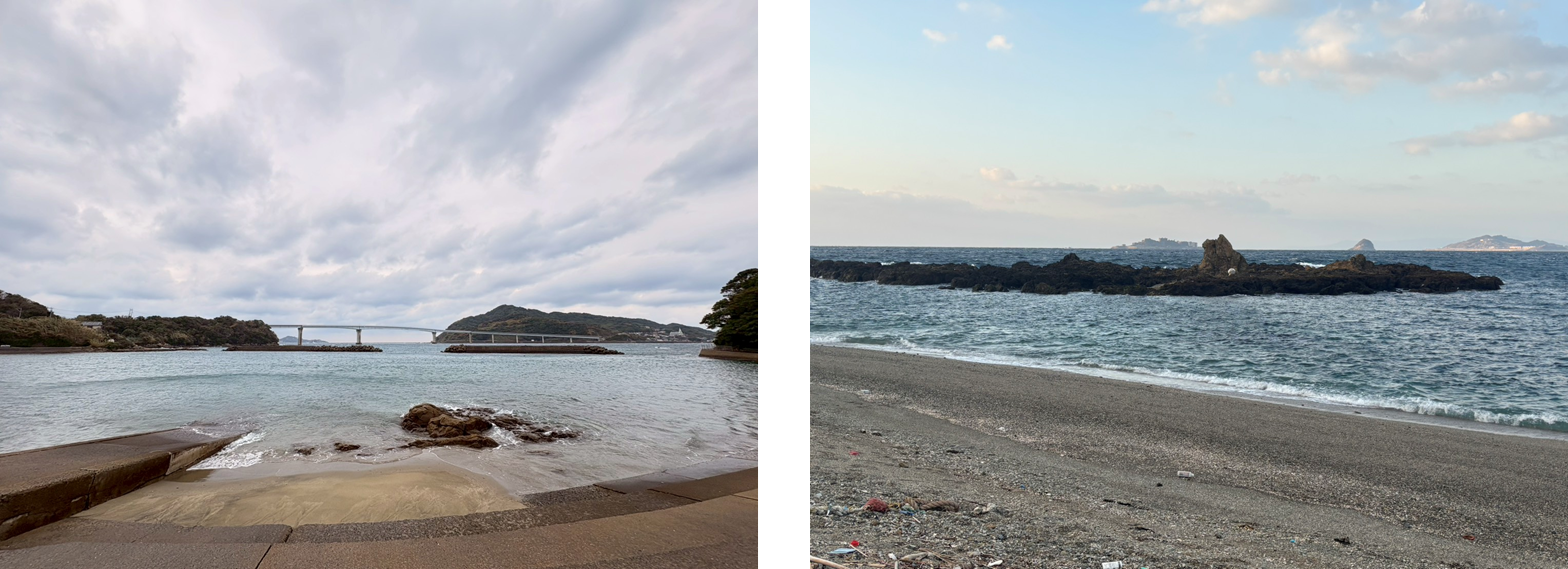

**Fig. S1** Photographs of the survey sites used in this study. Left: Tatsunokuchi; Right: Nomozaki Akase.

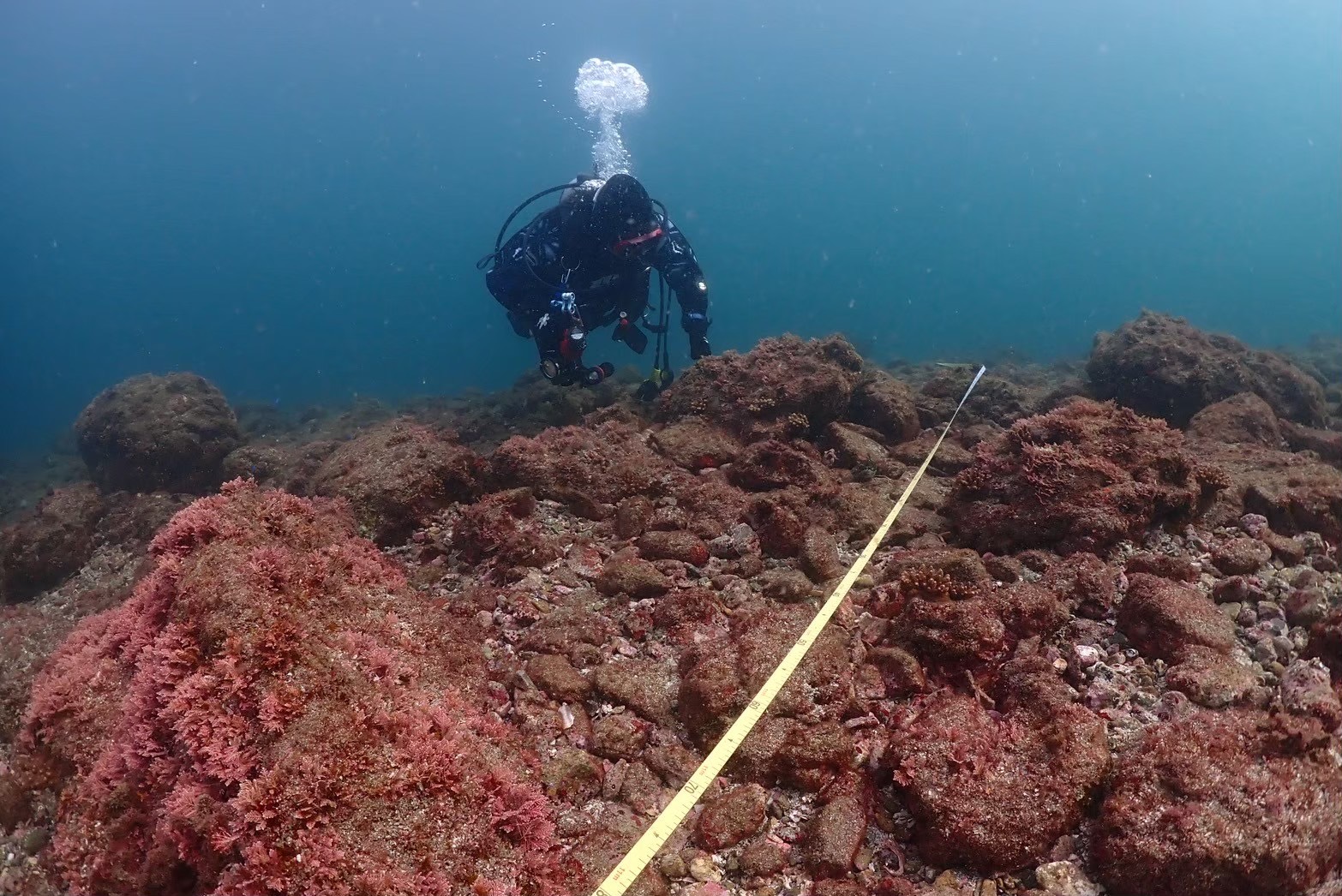

**Fig. S2 Photographs showing the diving surveys conducted in this study.**

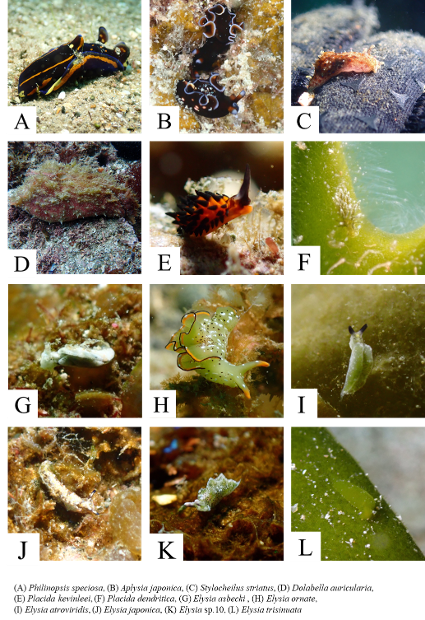

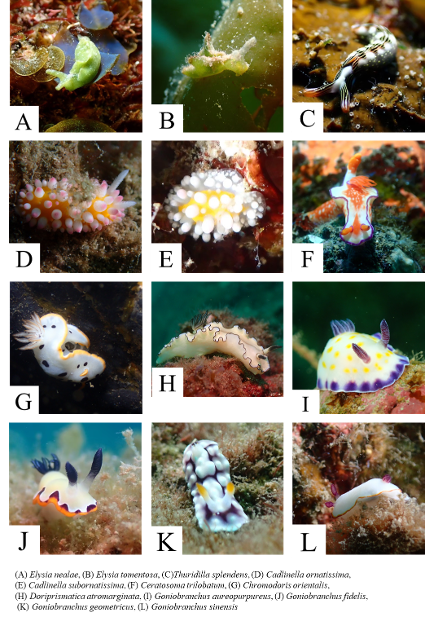

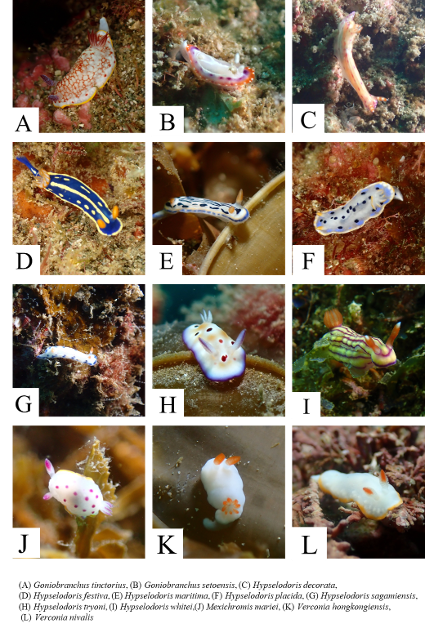

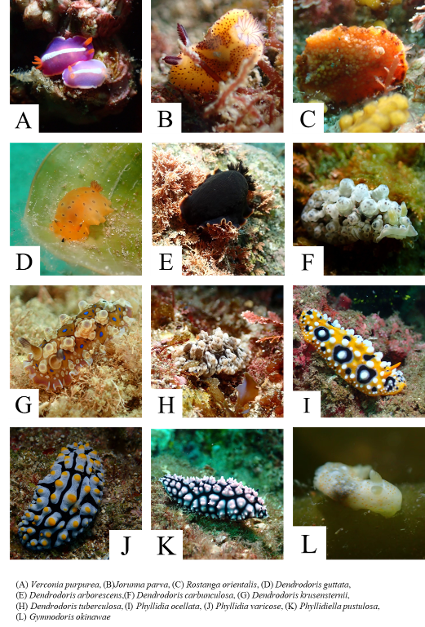

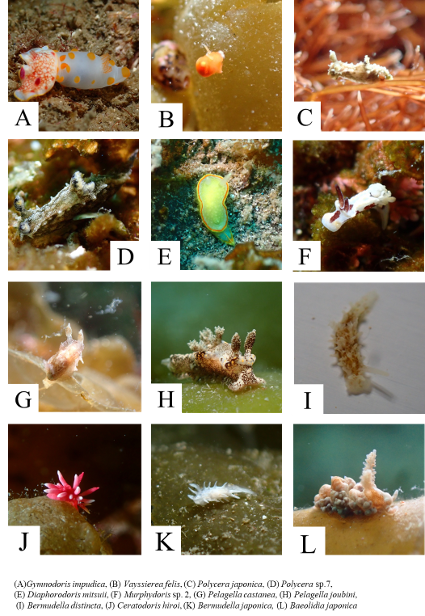

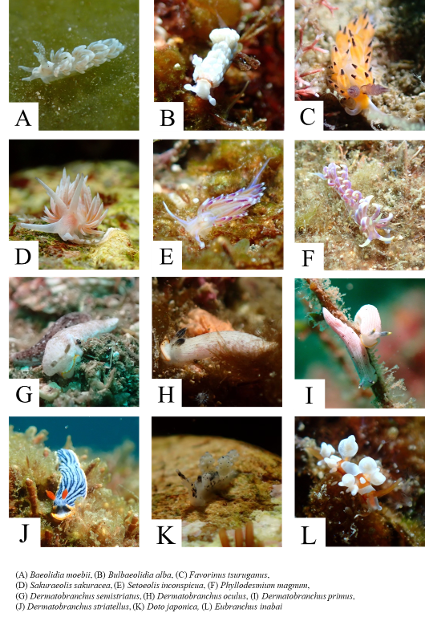

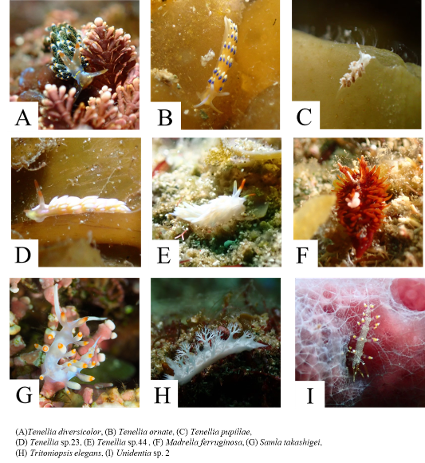

**Fig. S3** Photographs illustrate the morphological diversity of heterobranch sea slugs recorded from coastal waters of northwestern Kyushu, Japan, during the 2024–2025 underwater surveys.
